# Connecting spatial regions to clinical phenotypes by transferring knowledge from bulk gene expression data

**DOI:** 10.64898/2025.12.12.693322

**Authors:** Bayarbaatar Amgalan, Ermin Hodzic, M.G. Hirsch, Ziynet Nesibe Kesimoglu, John D. Bridgers, Jan Hoinka, David Levens, Chi-Ping Day, Teresa M. Przytycka

**Affiliations:** Computational Biology Branch, Division of Intramural Research, National Library of Medicine, NIH, Bethesda, Maryland, USA; Department of Computer Science, University of Maryland, College Park, Maryland, USA; Laboratory of Pathology, Center for Cancer Research, National Cancer Institute, NIH, Bethesda, Maryland, USA; Cancer Data Science Laboratory, Center for Cancer Research, National Cancer Institute, NIH, Bethesda, Maryland, USA

**Author notes:** co-first authors (Bayarbaatar Amgalan, Ermin Hodzic). SpacePhenotyper is implemented in Python. The source code and data sets used for and generated during this study are available at https://github.com/NLM-DIR/SpacePhenotyper.

## Abstract

Understanding intra-tumor heterogeneity is a key step towards understanding cancer. Spatially resolved transcriptomics (SRT) technology allows for estimation of transcriptional variability, cell types, and spatial relations between locations in a 2D tissue slice. In particular, it can be used to estimate molecular phenotypes of local regions, such as tumor purity and immune cell enrichment. In contrast, current methods remain limited in their ability to map clinical phenotypes to spatial locations, such as hazard or drug response. These clinical phenotypes are measured at the patient level and therefore require transferring knowledge from the patient level domain to the spot level domain. Here we present SpacePhenotyper, a new method that uses algebraic spectral techniques to transfer the predictive relationship between bulk gene expression and clinical phenotypes to the spatial domain. Our approach introduces the “Eigen-Patient,” a vector that is trained to predict the phenotype in bulk data and then used to quantify the phenotype in spatial spots. After validating SpacePhenotyper on both simulated and real data, we utilize it to study spatial heterogeneity of residual cancer burden in breast cancer treatment and and show that high proliferation alone does not predict treatment outcome. SpacePhenotyper has proven to be a powerful tool for transferring knowledge from bulk to spatial data.

## 1 Introduction

Intra-tumor heterogeneity is a challenging property of solid tumors with important consequences for cancer treatment. Many critical tumor properties, such as invasiveness or growth, depend on both specific transcriptomic changes in tumor cells and the tumor microenvironment (TME) — a complex, spatially organized ecosystem of diverse cell types surrounding tumor cells [1]. Interactions between tumor cells and the TME impact tumor growth and spread, therefore contributing to clinical outcomes [1, 2]. These complex relationships, however, are currently not fully understood.

Spatially resolved transcriptomics (SRT) technologies have enabled new insights into tumor heterogeneity [3, 4, 5, 6, 7]. These technologies provide spatial information on gene expression, typically allowing for an estimation of the transcriptional variability over locations in a 2D section of tissues. Popular platforms include those utilizing spotted arrays of mRNA-capturing probes on the surface of glass, capturing up to 10 cells per spot (e.g. 10X Genomics Visium), and *in-situ* hybridization of locally amplified gene-specific probes (e.g. 10X Genomics Xenium).

Until recently, analyses of SRT data have been mostly focused on identifying cell types present in spatial spots by adapting existing deconvolution methods based on cell-type-specific expression information from single-cell RNA sequencing [8, 9, 10, 11, 12, 13]. So inferred cell type information allows for studying tissue composition, predicting cell-cell [14, 15, 16, 17], and inferring certain *molecular phenotypes* of spatial regions, including the annotation of tumor versus stromal regions, regions with immune cell activity, and the identification of differentially expressed regions. Using these and other approaches, several methods to infer molecular properties of spatial regions have been recently developed. For example, Estimate [18] uses gene set variation analysis (GSVA) to quantify the presence of two molecular phenotypes of interest. It was initially developed for bulk gene expression analysis, and has recently been adopted to quantify the molecular activity of tumor and TME cells in spatial locations [19].

In contrast to estimation of molecular phenotypes, assignment of *quantitative clinical phenotypes* (traditionally measured on the patient level) to SRT regions has been left largely unaddressed, and direct application of classic supervised learning approaches has not been successfully applied to quantitative clinical phenotype prediction. Clinical phenotypes that are especially relevant for spatial data analysis include region-specific hazard or response to treatment, and their inference in spatial regions is critical for clinical studies due to their immediate relevance to treatment. A machine learning method CloudPred [20] was developed for binary patient classification from single cell data, but its binary classification design is not suitable for general clinical phenotype prediction. A study by Shulman *et al* [21] clustered localized spatial domains based on gene expression estimated from H&E images of a large patient cohort, and used them to stratify TCGA breast tumors based on spatial architecture and to predict patient-level responsiveness to treatment (as features for logistic regression). However, their study does not offer a method of assigning clinical phenotypes to spatial regions with intra-tumor heterogeneity. Notably, two recent methods leverage knowledge transfer from patients’ bulk data in their spatial data analysis. SpaLinker [22] decomposes bulk expression of phenotype-associated genes into non-negative “factors”, and subsequently infers their activities in spatial data. However, these factors are not necessarily interpretable in relation to the phenotype, and the associated genes’ expression is likely to carry additional phenotype-unrelated signal that ultimately gets reflected in the factors. SpaPheno [23] uses bulk cell type proportions as features from which to learn the patients’ phenotype quantities. The limitations of this approach include attributing the phenotype entirely to cell-type composition, thereby neglecting cell-type specific variation in gene expression, as well as relying on accurate inference of cell type proportions of both bulk samples and spatial locations.

In this paper, we introduce SpacePhenotyper, a method developed to overcome the challenges of predicting clinical phenotypes for spatial regions. We assume that a quantitative clinical phenotype of interest is predictable from gene expression both on the patient level (from bulk data), as well as on spatial region level (from gene expression of local spatial regions); see Discussion. SpacePhenotyper transfers such predictive relations from bulk data to the SRT domain. To that end, we introduce two related vectors termed *Eigen-Gene* and *Eigen-Patient*. The *Eigen-Gene* is a patient-wise vector optimized to predict relative patient-level phenotype from bulk gene expression data. The second vector, the *Eigen-Patient*, is a gene-wise vector that is derived from the *Eigen-Gene*, and is used to estimate the relative phenotype quantities from gene expression of spatial locations.

We evaluated our method on synthetic bulk and SRT data where spatial cell type landscape simulates tumor vs immune cell activity, and a molecular phenotype dependent on gene expression in certain cell subpopulations. We benchmarked SpacePhenotyper against other methods that utilize knowledge transfer from bulk to SRT data, such as SpaLinker and SpaPheno, as well as linear and non-linear regression models trained directly on bulk expression. None of the tested alternative approaches matched the accuracy of SpacePhenotyper, highlighting the limitations of factorization-based and cell type distribution based approaches.

Next, we validated SpacePhenotyper on real breast and colon cancer datasets using a set of pathologist-annotated H&E images and biological knowledge as standards of truth. Annotation of these images includes both tumor regions and tumor invasiveness, allowing us to infer the relationship between molecular phenotypes and *hazard* and *drug resistance* clinical phenotypes. For example, our analysis revealed that invasive tumor regions predicted to have high *hazard* are generally predicted to respond well to treatment. While these specific tumor regions also have high proliferation, our results show that high proliferation alone does not predict treatment outcomes.

## 2 Results

### 2.1 The SpacePhenotyper method

Given a bulk gene expression matrix *B* and phenotype information Φ for a group of patients, SpacePhenotyper infers two vectors that are associated with the phenotype, one in the gene space of *B*, and the other in the patient space (Fig.1). The first vector, the *Eigen-Gene*, is a patient-wise vector that is optimized to predict the phenotype, as a linear combination of principal directions of bulk expression in the gene space (Fig. 1A). This vector is then transformed into its image, the *Eigen-Patient*, which represents a predictor of the phenotype in the patient space (Fig. 1B). The coordinate values of the *Eigen-Gene* represent the predicted relative phenotype quantities across bulk patients, whereas the coordinate values of the *Eigen-Patient* represent the estimated relative predictive contributions of individual genes to the phenotype Φ.

**Figure 1:**
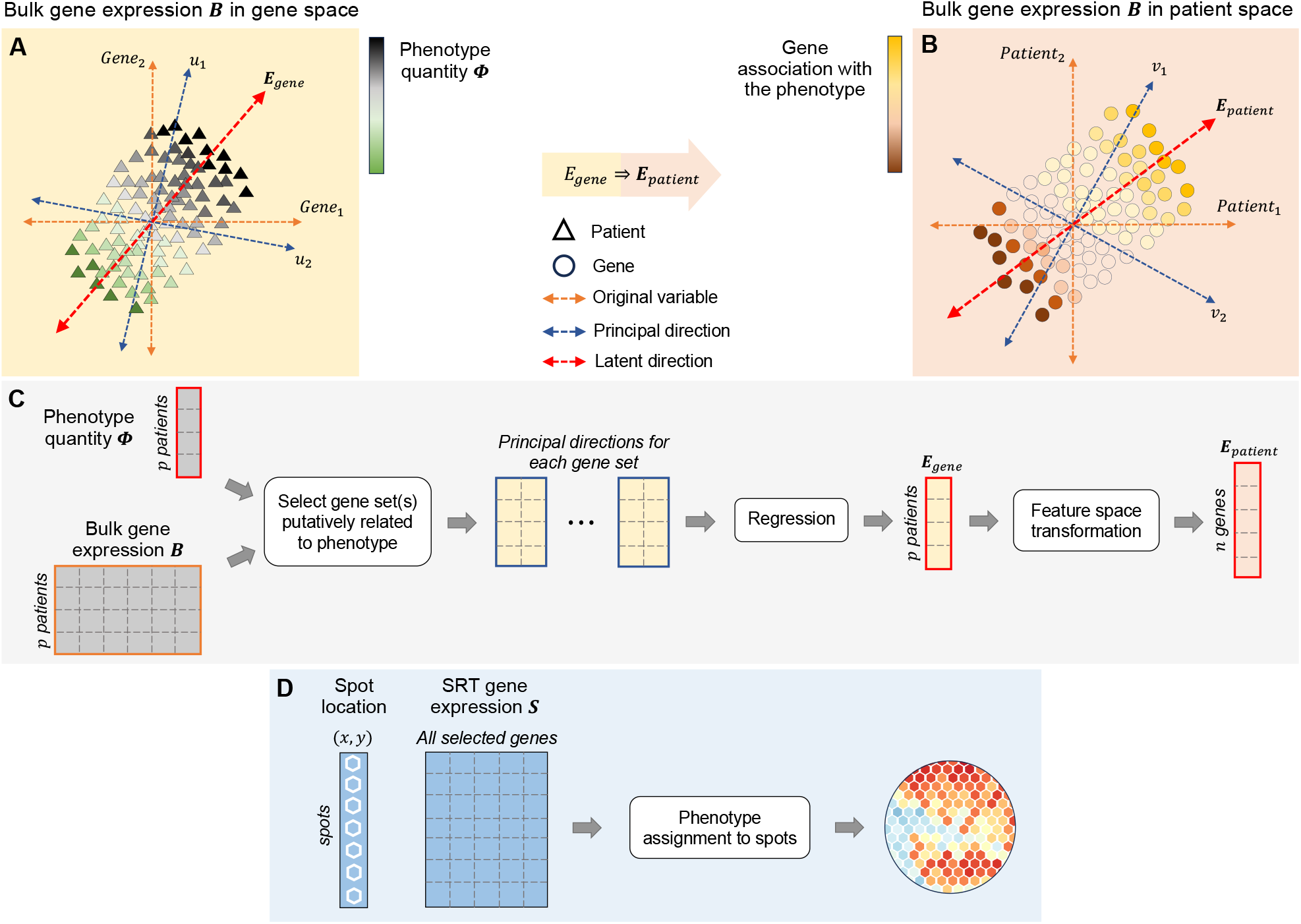
Workflow of the SpacePhenotyper method. **(A, B)** Conceptual overview of transforming a vector modeling relative phenotype quantity in gene space to its image in patient space. The transformation takes *Eigen-Gene*, a vector of values across patients, as input and produces the corresponding *Eigen-Patient*, a vector of values across genes, as output. **(C)** SpacePhenotyper workflow. SpacePhenotyper takes three inputs: the phenotype quantity Φ, the corresponding bulk gene expression data *B* (gray), and spatially resolved transcriptomic data *S* (blue). For the given phenotype, it creates the *Eigen-Gene*, a gene-space predictor of the phenotype, and its corresponding patient-space image *Eigen-Patient*. **(D)** *Eigen-Patient* is used as the predictor to assign relative quantity of the phenotype in each spot (*x*_*i*_, *y*_*i*_).

#### 2.1.1 Learning the phenotype from expression of disjoint gene sets

Let *B* _*p*× *n*_ be the bulk expression matrix for *p* patients and *n* genes. According to the *Singular Value Decomposition (SVD)* theorem, the compact SVD of *B* is the factorization of *B* into *U* × Σ × *V*^*T*^, where Σ_*r*× *r*_ (*r* ≤ min {*n, p*}) is a diagonal matrix with elements *σ*_1_ > *σ*_2_ > … > *σ*_*r*_, which are positive singular values uniquely determined by *B*. The matrices *U*_*p*× *r*_ and *V*_*n*× *r*_ are semi-orthogonal, with columns denoting the left singular vectors *u*_*i*_ ( *i* = 1, …, *r*) and right singular vectors *v*_*i*_ (*i* = 1, …, *r*) of *B*, respectively. The left singular vectors represent the principal directions of *B* in its gene space, whereas the right singular vectors represent the principal directions of *B* in its patient space (Fig. 1A and B). From *B* = *U*Σ*V*^*T*^ and the semi-orthogonality of *U* and *V*, we obtain *U* = *BV*Σ^−1^ and *V* = *B*^*T*^*U*Σ^−1^. Consequently, for each pair of corresponding singular vectors *u*_*i*_ and *v*_*i*_, *u*_*i*_ = *Bv*_*i*_*σ*_*i*_ ^−1^, and *v*_*i*_ = *B*^*T*^*u*_*i*_*σ*_*i*_ ^−1^. More generally, let *I* be a subset of indices from 1 to *r*, and let *U*_*I*_, *V*_*I*_ and Σ_*I*_ denote the matrices obtained by retaining only the singular vectors and singular values indexed by *I*. Then, 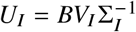 and 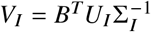.

The first principal direction captures the greatest amount of variation in the bulk gene expression matrix *B*. Each subsequent principal direction captures progressively less variation while remaining orthogonal to all preceding directions. In the remainder of methods, we will use the notation *U*^[*k*]^, Σ^[*k*]^ and *V* ^[*k*]^ to denote the SVD matrices obtained by retaining only the first *k* principal directions. The number of principal directions, *k*, will be treated here as a fixed parameter, with its optimal value determined through cross-validation, as described later.

Let Φ be a vector of patient-wise continuous phenotype quantities, and let *G*_1_, …, *G*_*m*_ be a collection of *m* mutually disjoint sets of genes. Each set *G*_*i*_ may represent a distinct molecular process or other biological property associated with Φ, with potentially different relationships to the phenotype. Here, we describe how to construct a predictor of Φ by integrating the predictive information contributed by each gene set *G*_*i*_. For 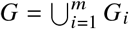, let *B*_*G*_ denote the submatrix of *B* containing only the columns corresponding to genes in *G*. For each gene set *G*_*i*_, we construct a matrix *B*_*i*_ from *B*_*G*_ by retaining the columns corresponding to genes in *G*_*i*_ and setting all other columns to zero. Each resulting matrix is then decomposed independently using singular value decomposition (SVD): *B*_*i*_ = *U*_*i*_Σ_*i*_*V*_*i*_^*T*^ . Given a selection of the first *k*_*i*_ principal directions from each *B*_*i*_, we model the phenotype as follows:

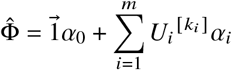

where *α*_*i*_ denote vectors of coefficients for the gene sets’ principal directions. The optimal coefficient vectors *α*_1_, …, *α*_*m*_ and the intercept *α*_0_ are then computed by solving the following least-squares optimization problem, which minimizes the estimation error measured by the squared Frobenius norm:

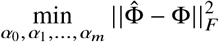

We next show that 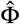 is a linear function of gene expression. From the SVD of each *B*_*i*_, we have 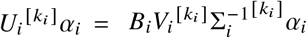. Since *B*_*i*_ has zero-valued columns corresponding to genes outside *G*_*i*_, and because 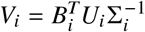, the rows of *V*_*i*_ (including those of 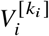), corresponding to genes outside *G*_*i*_ are also zero. Consequently, it follows that 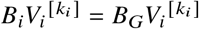. Then:

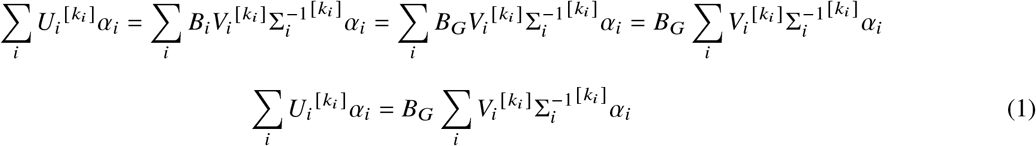

The left-hand side of Equation (1) represents the optimal linear combination of the selected principal directions associated with the disjoint gene sets that best predicts the relative phenotype quantities Φ, excluding the intercept. We refer to this vector as the Eigen-Gene. Equation (1) shows that the predicted relative phenotype quantities can be expressed as the dot product of the gene expression matrix *B*_*G*_ and a column vector, 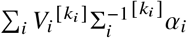 which serves as a linear predictor of the relative phenotype based on gene expression. This vector is itself a linear combination of the corresponding right singular vectors associated with the individual gene sets. We refer to it as the *Eigen-Patient*. Thus, the *Eigen-Gene* and *Eigen-Patient* are related through the linear transformations defined by *B*_*G*_ and its pseudoinverse, respectively: *E*_*gene*_ = *B*_*G*_ × *E* _*patient*_ and 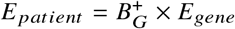, where 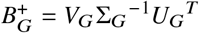 is the Moore–Penrose pseudoinverse of *B*_*G*_. Therefore, the predicted phenotype can be equivalently expressed as

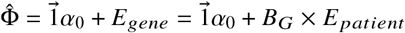

Finally, consider a spatial location (*x*_*i*_, *y*_*i*_) with a gene expression profile *s*_*i*_. Its relative phenotype quantity, *ϕ* (*x*_*i*_, *y*_*i*_), can be predicted by taking the inner product of its gene expression profile and the *Eigen-Patient*. Specifically, let *s*_*i*_ (*G*) denote the vector of expression values for genes in *G*, arranged in exactly the same gene order as the columns of *B*_*G*_. The predicted relative phenotype quantity at this spatial location is then

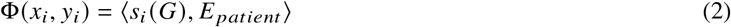

#### 2.1.2 Learning relative *hazard*

For *hazard* analysis, rather than being given a phenotype vector Φ containing the phenotype quantity for each patient, we are given a pair of vectors Φ = (*δ, t*) ; where *δ* denotes the patients’ vital status, and *t* denotes the time-to-event outcome. Specifically, for the *i*-th patient, *t*_*i*_ represents the survival time if *δ*_*i*_ = 0 and the censoring time if *δ*_*i*_ = 1. We then employ the Cox proportional hazards model [24] to model the patients’ hazard, using the principal directions in the gene space as covariates:

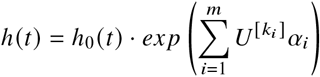

Here, the hazard function *h*(*t*) is determined by the covariates, with their respective effects quantified by the coefficient vectors *α*_*i*_. The term *h*_0_(*t*), referred to as the baseline hazard, denoting an intercept term that varies with time. Accordingly,

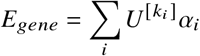

represents the estimation of relative log-hazard for bulk patients, and *ϕ* (*x*_*i*_, *y*_*i*_) = ⟨*s*_*i*_ (*G, E*) _*patient*_⟩ provides a gene-expression-based estimate of the relative log-hazard at each spatial location.

#### 2.1.3 Selection of gene sets and principal directions

SpacePhenotyper uses two gene sets for each phenotype: genes that are positively associated with the phenotype, and genes that are negatively associated with the phenotype. Specifically, for continuous phenotypes we calculate the association *a*_*i*_ of each gene *g*_*i*_ as the coefficient of Pearson correlation between its gene expression and the phenotype. For the *hazard* phenotype, we calculate *a*_*i*_ as the coefficient of univariate Cox regression in which its vector-length-normalized gene expression is the covariate. We then select the top 10% of genes1 with the largest non-zero absolute value of association coefficients |*a*_*i*_| to form the set *G*, which is subsequently partitioned into two disjoint subsets: *G*_1_, containing genes with *a*_*i*_ > 0, and *G*_2_ containing genes with *a*_*i*_ < 0.

We determine the number of top principal directions to retain for the two gene sets in Equation (1) as follows. We identify the smallest integer *k*_*max*_ such that in either of the two sets it holds that the first *k*_*max*_ principal directions explain at least 99% of the total gene expression variance captured by all of the set’s principal directions. If either gene set is empty (i.e. either all genes in *G* are positively associated with Φ, or all of them are negatively associated with Φ), that set is excluded from this calculation. The proportion of variance explained by the first *k* principal directions is calculated as

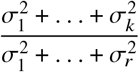

After determining *k*_*max*_, we train Eigen-Patient for each for each *i* = 1, …, *k*_*max*_, setting *k*_1_ = *k*_2_ = *i* in Equation (1). Model training and evaluation are performed using 5-fold cross-validation, with 100 random permutations of the sample order before fold assignment. We select the *Eigen-Patient* with the highest average cross-validation (CV) score as the final model returned by SpacePhenotyper. The CV score is measured using the Pearson correlation coefficient between predicted and observed phenotype values for continuous phenotypes, and the concordance index (C-index) for *Hazard*.

### 2.2 SpacePhenotyper accurately predicts phenotypes of spatial regions in simulated data

We begin by validating SpacePhenotyper’s accuracy of phenotype prediction on simulated data. A comprehensive analysis of clinical phenotypes in tumor slices also necessitates prediction of molecular phenotypes, allowing for a deeper investigation into the relation between molecular mechanisms and higher level phenotypes. Since all tested methods are agnostic to the type of phenotype, here we simulate data for molecular phenotypes.

Synthetic data was generated to reflect the following basic assumptions: i) patients’ phenotype quantities are predictable from their bulk expression, ii) the predictive relation learned in the bulk domain can be transferred to the spatial domain, and that iii) the simulated phenotype is associated positively with gene expression in a specific cell subpopulation and negatively in another. Given the above, we generated SRT data with four spatial regions with distinct distributions of epithelial, immune, and stromal cells (Fig. 2A), and each cell’s gene expression was sampled from real single-cell data. The region labeled “Neutral” contains only stromal cells, the one labeled “A” contains almost entirely epithelial cells, the one labeled “B” contains almost entirely immune cells, and the region labeled “Mixed” contains a mix of all three cell types. We select two subsets of genes that are shared by all three cell types, with the gene expression of one set contributing positively to the phenotype in those epithelial cells in which they are highly expressed, and the gene expression of the other set contributing negatively to the phenotype in all immune cells. Ground truth phenotype quantities in every spot were computed as a function of these positive and negative contributions by each cell in that spot (Fig. 2B, C), resulting in a nonlinear relationship between the phenotype, gene expression, and cell type distribution. Gene expression and phenotype quantities for bulk data patients were computed analogously. For the exact details of synthetic data generation, see Supplementary Materials Section S-1.2.

**Figure 2:**
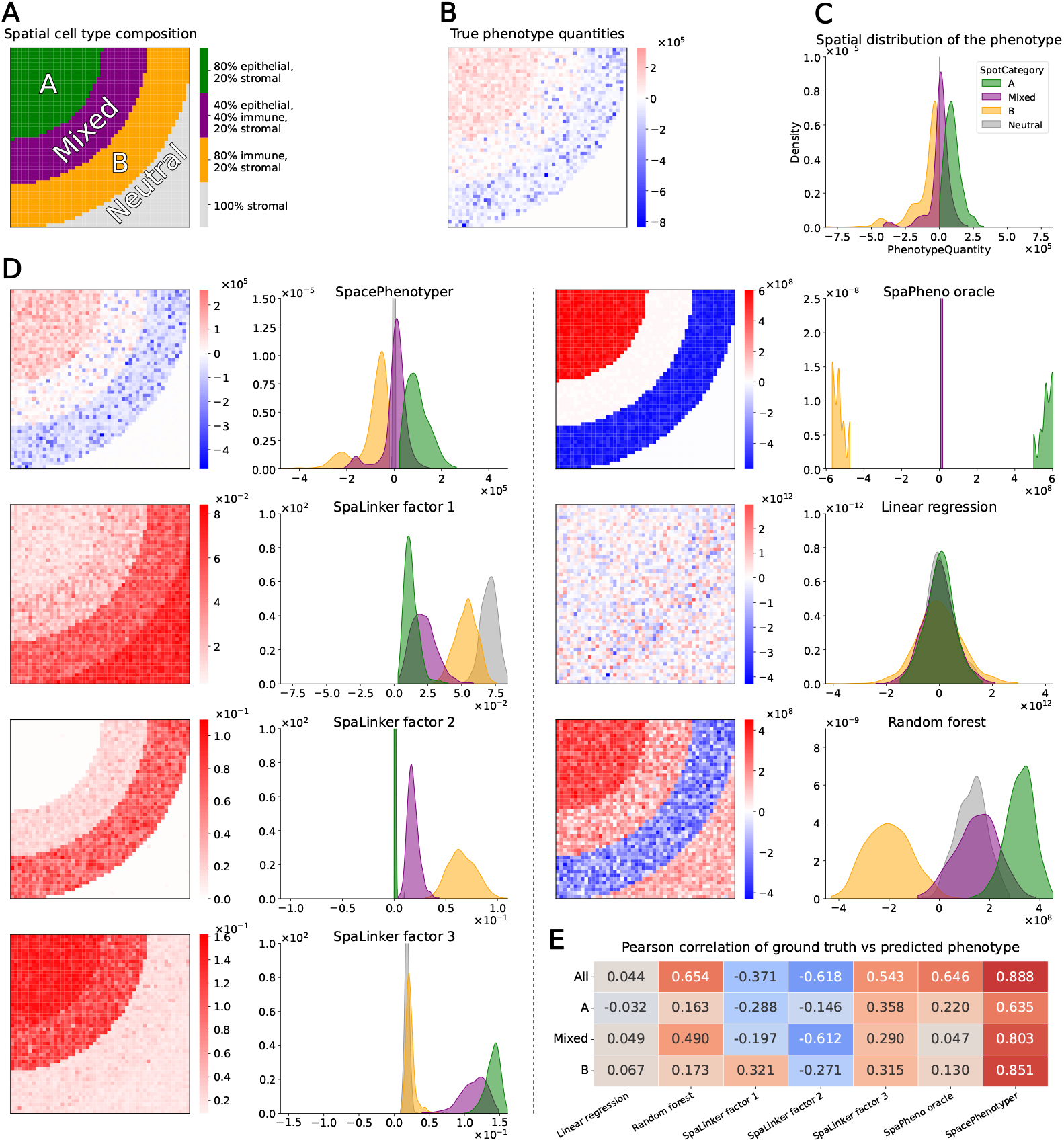
Evaluation of phenotype prediction on synthetic data. Synthetic pseudo-bulk and pseudo-SRT data was generated by sampling single-cell expression of three cell types from a real dataset. The quantity of the phenotype was simulated to depend on gene expression in a manner that is linked to, but not entirely dependent on, cellular composition. Please see supplementary methods for a detailed description of the simulation procedure. The resulting bulk expression, SRT expression, and bulk patient phenotype quantities were given as input to the tested methods. Note that ground truth cell type proportions for each spot were given to SpaPheno, to provide it with the best conditions, and we name it “SpaPheno oracle” in the results. **(A)** Outlines of 4 synthetic spatial regions with distinct cell type compositions on a 50x50 slice. **(B)** Heatmap showing true quantities of the simulated phenotype in each spatial location. **(C)** Distribution of true phenotype quantities across the 4 distinct spatial regions. **(D)** For each tested method, a heatmap of predicted phenotype quantities in SRT spots, and the corresponding distribution of predicted phenotype quantities across the 4 regions. For SpaLinker, all returned factors are displayed. **(E)** Pearson correlation coefficients for ground truth phenotype quantities vs predicted values; both across the entire slice, as well as individual regions.

We tested SpacePhenotyper’s performance against SpaLinker and SpaPheno, two existing methods that also leverage knowledge transfer from bulk data. Additionally, we tested alternative baseline approaches such as a direct application of linear and random forest based regression on the bulk gene expression data. All tested methods were given the bulk gene expression, bulk patient phenotype quantities, and SRT gene expression data as input. Notably, the method SpaPheno requires additional inputs in the form of estimated cell type proportions for each patient and each spatial location, and we provide it with the most advantageous setting by giving it ground truth cell type proportions — in the results, we label this advantaged version “SpaPheno oracle”. We also note that we disabled its SRT expression smoothing (unnecessary for this benchmark), which is enabled by default.

The results shown on Fig. 2(D, E) demonstrate the high accuracy of SpacePhenotyper at correctly predicting the simulated phenotype via all measured metrics: visually, both the heatmap and the density of its predicted phenotype quantities resembles the ground truth most accurately, and its predictions achieve the highest Pearson correlation coefficient with the ground truth across all individual regions, as well as the entire slice (0.888). SpaPheno found the correct outlines of the four spatial regions and their overall positive (region “A”) and negative (region “B”) association with the phenotype, but was unable to capture differences in phenotype quantities within those regions, which is perhaps most evident in the “Mixed” region where it estimated the phenotype to be near zero. SpaLinker, which returned three factors, also showed the ability to capture region outlines, but none of the factors capture the phenotype itself on their own. We speculate that both these methods’ phenotype estimation accuracy might be confounded by the cell type distribution signal: in the case of SpaPheno directly from the given cell type distributions, and in the case of SpaLinker through expression of genes that are exclusive to one cell type. The random forest approach scored the second highest Pearson correlation coefficient for the entire slice (0.654), but it wrongly predicted the association with the phenotype in the “Neutral” region to be positive, did not closely resemble the phenotype quantities within other regions, and failed to identify split association with the phenotype in the “Mixed” region. The naive linear regression approach did not yield any accurate results. Overall, SpacePhenotyper clearly outperformed other approaches under the simulated conditions, and displayed outstanding results.

### 2.3 Validation of bulk to spatial knowledge transfer in biological data

In this section, we show that SpacePhenotyper can accurately predict phenotypes in real biological data. We apply our approach to predict two clinical phenotypes: *hazard* and *drug resistance* measured by Residual Cancer Burden (RCB) [25], as well as several molecular phenotypes including *tumor purity, immune response*, and *proliferation*, to allow for the discovery of associations between clinical and molecular phenotypes.

#### Bulk Data Sources

For tumor purity, the estimation of the *Eigen-Patient* is based on the results of [26] which provides tumor purity scores obtained by integrating multiple measurements including gene expression, DNA methylation, immunohistochemistry, somatic gene copy numbers, and image analysis. For the *proliferative* phenotype and the *immune* phenotypes, we performed single-sample gene set enrichment analysis (ssGSEA) [27], utilizing respectively the gene set involved in proliferation according to Gene Ontology (GO:0008283) and lymphocyte activation involved in immune response (GO:0002366). In these two cases, we ultimately use patients’ scores for enrichment in the respective gene sets as quantitative phenotype. For *hazard*, we use survival data from TCGA. For *drug resistance* we use the post-treatment residual cancer burden from TransNEO data [28].

#### Spatial Data Sources

In this study, we used several samples from Breast cancer including Breast 1.1.0 (in [13, 29, 30]), Breast 1.3.0 (in [31]), and Breast 2.0.0 (in [32]), and one Colon cancer sample Colorectal 3.0.0 (https://www.10xgenomics.com/datasets/visium-hd-cytassist-gene-expression-libraries-of-human-crc). Breast 1.1.0 and Breast 1.3.0 were obtained with Visium technology, 2.0.0 was obtained with Xenium, and Colorectal 3.0.0 was obtained with HD Visium. For details about the data its processing and normalization see Section 1.1 in Supplementary Materials.

#### 2.3.1 *Eigen-Patient* accurately predicts relative phenotype quantities in bulk expression

To assess the accuracy of SpacePhenotyper’s predictions in biological data, we performed cross-validation of the *Eigen-Patient* on the bulk patient data. The order of bulk samples was randomly shuffled 100 times, and 5-fold cross-validation was performed for each random order, yielding 500 paired training and test sample sets. For each pair, *Eigen-Patient* was computed from training samples, and used to predict phenotype quantities for the test samples. For continuous phenotypes, predictions were scored by their coefficient of Pearson correlation with the phenotype. For the *Hazard* phenotype, they were scored by the concordance index. Table 1 shows the mean prediction score over all 500 experiments, together with standard deviation in parentheses.

**Table 1:** Cross-validation of the Eigen-Patient. The order of bulk samples was randomly shuffled 100 times, and 5-fold cross-validation was performed for each random order, yielding 500 paired training and test sample sets. For each pair, *Eigen-Patient* was computed from training samples, and used to predict phenotype quantities for the test samples. For continuous phenotypes, predictions were scored by their coefficient of Pearson correlation with the phenotype. For the *Hazard* phenotype, they were scored by the concordance index. The table shows the mean prediction score over all 500 experiments, together with standard deviation in parentheses.

|  | Tumor purity | Proliferation | Hazard | Lymphocyte activation | Drug resistance |
|---|---|---|---|---|---|
| Breast 1.1.0 | 0.718 (0.035) | 0.905 (0.011) | 0.572 (0.062) | 0.799 (0.024) | 0.541 (0.109) |
| Breast 1.3.0 | 0.717 (0.036) | 0.905 (0.011) | 0.572 (0.062) | 0.799 (0.024) | 0.541 (0.109) |
| Breast 2.0.0 | 0.693 (0.044) | 0.904 (0.011) | 0.586 (0.060) | 0.808 (0.022) | 0.533 (0.105) |
| Colorectal 3.0.0 | 0.723 (0.073) | 0.902 (0.023) | 0.552 (0.100) | 0.588 (0.070) | N/A |

#### 2.3.2 Validation of clinical and molecular phenotype predictions in spatial regions in biological data

To validate SpacePhenotyper’s output from biological data, we scrutinized the consistency of predictions both with pathologist-annotated Hematoxylin and Eosin (H&E) staining images, as well as with current knowledge. The H&E images of breast cancer samples were annotated by pathologists to differentiate between various stages of tumor regions. Ductal carcinoma begins in the lining of the milk ducts as ductal carcinoma in-situ (DCIS), which is assumed to be a non-invasive tumor that is contained within the milk duct. Some DCIS gradually evolve into DCIS with microinvasion, where tumor cells begin to invade the tissue surrounding the milk ducts. Eventually, the tumor progresses to invasive ductal carcinoma (IDC) that spreads beyond the milk ducts (Fig. 3), and has the potential to metastasize. These different stages of tumor progression can co-exist in one sample. Pathologists can distinguish between DCIS, DCIS with microinvasion and IDC in H&E images based on the organization of the epithelial tissue surrounding the duct (Fig. 4A). In addition, a recent study identified a novel invasive-like DCIS — an in-situ tumor that expresses an invasive molecular signature. Breast 1.3.0, in addition to invasive and in-situ regions, also includes in-situ regions with potential microinvasion. There is no information on potential invasive-like in-situ regions for Breast 1.1.0 and Breast 1.3.0. Finally, Colorectal 3.0.0, in addition to tumor regions, has an annotated region of non malignant epithelial cells. Small regions which could be significantly influenced by boundary spots are omitted. The diversity of tumor stages in the annotated H&E images included in this study provides the opportunity to test whether SpacePhenotyper can predict phenotypic properties that match pathologist annotations and biological knowledge. Our results, discussed below, are visualized in Fig. 4 and quantified in Fig. 5.

**Figure 3:**
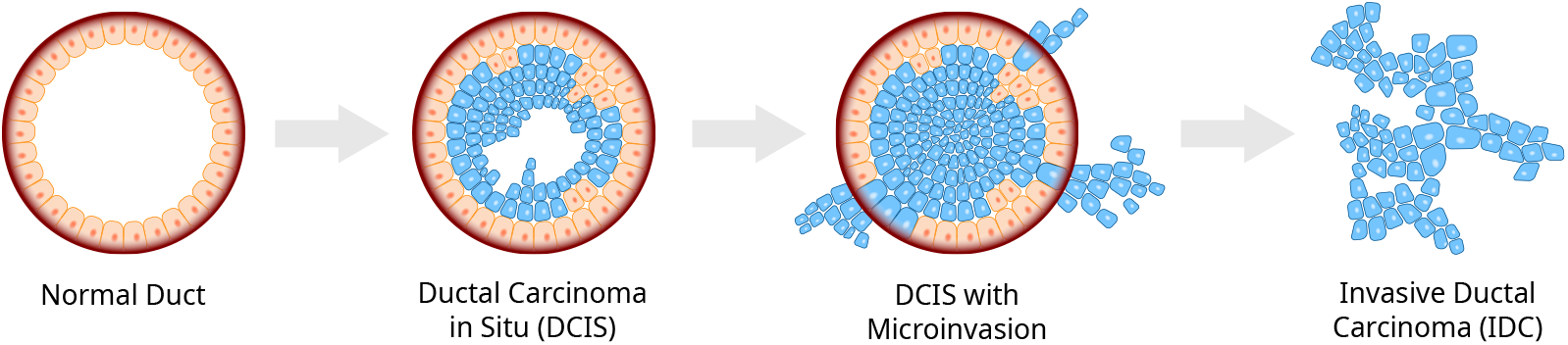
Stages of ductal carcinoma. Ductal carcinoma begins as ductal carcinoma in-situ (DCIS) in the lining of the milk. Some DCISs gradually evolve into DCISs with microinvasion, where tumor cells begin to invade the tissue surrounding the milk ducts. Eventually, the tumor progresses to invasive ductal carcinoma (IDC) that spreads beyond the milk ducts.

**Figure 4:**
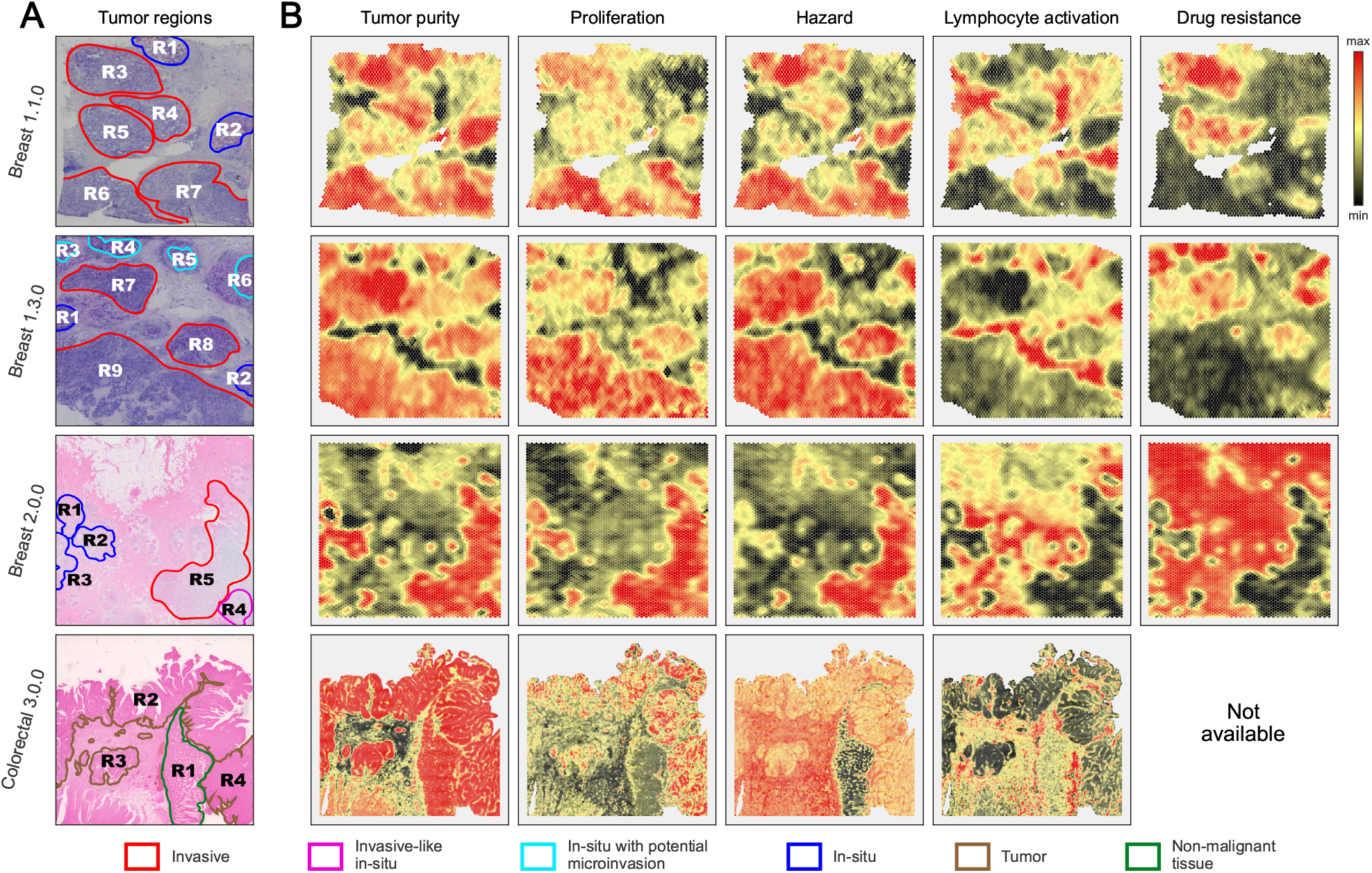
Spatial heterogeneity of phenotypes on SRT samples. Predictions of *tumor purity, lymphocyte activation* and *proliferative* molecular phenotypes, as well as two *c*linical patient phenotypes, drug resistance and *hazard*, on three BRCA and one COAD SRT sample. **(A)** Annotated H&E images showing outlines of different spatial tumor regions of interest. **(B)** Heatmaps for relative phenotype quantities predicted by SpacePhenotyper. Each heatmap’s color scale is independent.

**Figure 5:**
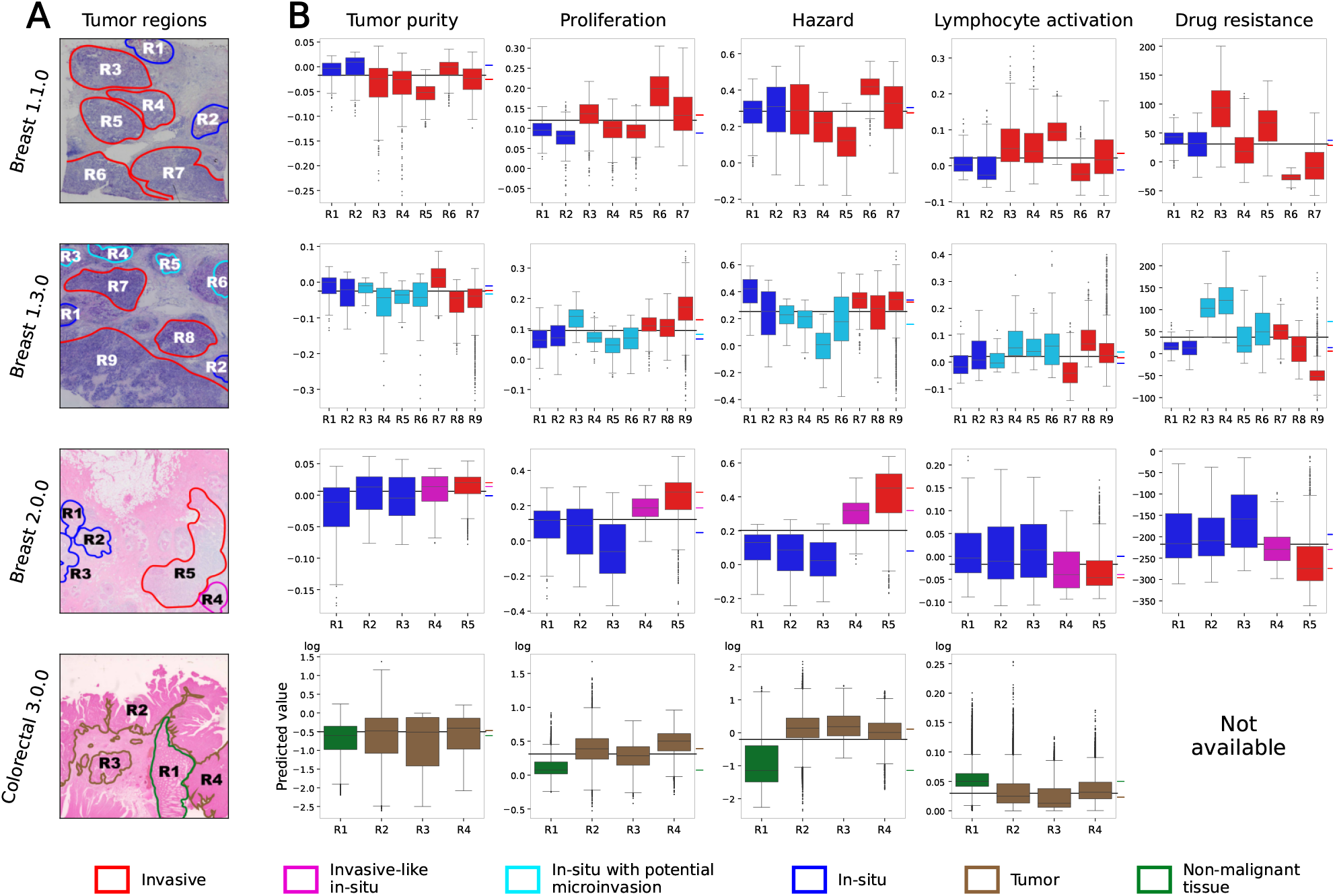
Phenotype distribution across spatial regions of interest. **(A)** Annotated H&E images showing outlines of different spatial tumor regions of interest. **(B)** Box plots summarizing the distribution of predicted phenotype values in each spatial region of interest .

##### Tumor purity and proliferation

First, we used pathologist-annotated H&E images (Fig. 4 A) as a reference to test whether annotated tumor regions are aligned with predicted high tumor purity and proliferative regions. Fig. 4 shows good agreement between predicted high *tumor purity* and pathologist-annotated tumor regions. As expected, in-situ tumor regions (dark blue in Fig. 4A, Fig. 5) and all but one in-situ with potential microinvesions (light blue) have lower proliferation than invasive tumors (red). In Colorectal 3.0.0, both tumor and non-malignant regions are predicted as high tumor purity, but the non-malignant tissue is less proliferative. The benign status of this region is consistent with the hazard analysis below.

##### Hazard

For the colorectal cancer sample, we leveraged a pathologist-annotated non-malignant region and confirmed that this region indeed exhibited low predicted hazard (Fig. 4 and Fig. 5). For Breast Cancer, we do not have a similar absolute gold standard of truth to rank hazard. However, Ru *et al*. [13] showed that regions R6 and R7, which are predicted to have the highest hazard in Breast 1.1.0, are enriched for a gene-expression signature associated with the complete mesenchymal-epithelial transition (cEMT) state, consistent with our prediction. In addition, we noted that a common property of all high hazard regions (R6, R7 on slide Beast 1.1.0; R1, R7, R9 side Breast 1.3.0, R4, R5 slide Breast 2.0.0 and R2, R3, R4 slide Colorectar 3.0.0) is a relatively high predicted lymphocyte activation (Fig. 5), while low hazard regions show a reverse property. This is consistent with biological understanding that lymphocyte activation emits adaptive and innate-like immune response. Interestingly, the in-situ tumor regions were sometimes predicted to be more hazardous than some invasive tumors. All in-situ tumors are often considered to be non-invasive. However, as shown in [13], some in-situ tumors are expressing an invasive molecular signature. In-situ region R4 in Breast 1.3.0 has such invasive signatures. Consistent with its invasive signature, our prediction shows that this region has high hazard.

##### Applicability of SpacePhenotyper to HD Visium data

The applicability of SpacePhenotyper to the HD Visium panel (Colorectal-3.0.0 sample) was not immediately evident, as in this High Definition setting one spot can harbor a fragment of a cell rather than one or multiple cells. While the phenotypic estimation of spots was somewhat noisy for Colorectar 3.0.0 sample (Fig. 5y axes), SpacePhenotyper correctly identified phenotypic regions of this slide, including the proper recognition of the non-malignant tissue region as high purity and relatively low proliferativity and not associating it with hazard (Fig. 4). In addition, SpacePhenotyper’s predicted high hazard of stroma regions is consistent with the observation that tumor stroma percentage (TSP) is a marker of poor prognosis in colorectal cancer [33].

### 2.4 Analysis of drug response prediction

Encouraged by the success of SpacePhenotyper for other phenotypes, we applied it to predict regional drug response. In treatment of breast cancer, a course of neoadjuvant chemotherapy is often administered prior to a potential surgery to shrink the tumor. Residual cancer burden (RCB) is used to quantify the response to neoadjuvant treatment [25]. These treatments are often targeted towards the invasive tumor regions and not the in-situ tumor regions (stage 0). However, given that some of these in-situ tumors might carry invasive molecular signatures, like R3-R6 in Sample 1.3.0 and R4 in Sample 2.0.0, it is important to know whether such regions would respond to the therapy. If so, neoadjuvant chemotherapy for earlier stages might be desirable.

With the goal of identifying tumor regions that would respond to neoadjuvant therapy, we trained our predictor on the TransNEO data [28] containing bulk expression and RCB phenotype quantities. Previous studies used this data to construct a predictor of patient-level phenotype using spatial clusters of gene expression as features, but did not analyze the drug response of different tumor regions. In contrast, we focus on predicting the drug response of the latter, allowing us to explore the intra-tumor heterogeneity of this response and the dependency of this response on molecular phenotypes of spatial tumor regions.

Our results suggest that regions that have high proliferation or hazard generally respond well to the treatment. There was only one major exceptions: R3 in Sample 1.3.0 which is an in-situ tumor with potential microinvasion. This region has high drug resistance despite high proliferation. In addition, the response of region R4 (invasive-like in-situ) is sample 2.0.0 is modest. This suggests that treating proliferative tumors while still contained to the milk duct may not be effective. Interestingly, both regions show low lymphocyte activation in line with the observation that higher levels of tumor-infiltrating lymphocytes is associated with a higher rate of complete response [34]. This suggests that high proliferation and hazard are good biological phenotypes to predict good response, but activated lymphocytes also play a role. We also observed that stroma in sample 2.0.0 expresses drug resistance signature. Stroma has no proliferative phenotype so the drug is not expected to have an effect on this tissue. This might have lead to drug resistance-like signal and underscores the importance of considering the prediction in the context of other molecular phenotypes which jointly define the properties of a spatial region.

Our results suggest heterogeneity of responses in different regions of the same tumor. In addition, although the proliferating cells are the primary targets of this therapy, high proliferation alone does not guarantee a good outcome. It would be important to confirm these emerging patterns in a larger cohort.

## 3 Discussion

Due to intra-tumor heterogeneity, different tumor regions are likely to have different phenotypic properties. Inferring clinical phenotypes of spatial regions is a first step towards understanding of clinical consequences of this variability. A formidable obstacle to this inference stems from the fact that clinical phenotypes, such as hazard or therapeutic response, are measured on the patient level rather than on the level of spots or other spatial tumor regions. Therefore, to assign clinical phenotypes to spatial regions, one needs to transfer knowledge from patient level data to spatial data. We design SpacePhenotyper to address this task.

The key computational concept introduced in this work is the *Eigen-Patient* — a predictive vector of gene weights, constructed in a way that enables the transfer of knowledge between the patient and spatial domains. The construction of *Eigen-Patient* utilizes principal directions. However, rather than being based on all genes, it is computed based on principal directions of gene groups, allowing it to account for diversity of responses to phenotypic variations in different gene groups. Our analysis was performed utilizing two groups of positively and negatively correlated genes, but we also give a general theoretical framework for an arbitrary number of disjoint gene groups. While in this work we define *Eigen-Patient* as a linear predictor, our theoretical framework can be generalized to non-linear predictors. Our tests show the robustness of our approach even when the number of samples from which to learn a phenotype predictor is not large, a scenario that would be challenging for more complex deep learning approaches.

SpacePhenotyper was developed under the assumption that a local phenotype depends on (more precisely is predictable by) spatially local gene expression. This assumption is clearly true for many phenotypes including tumor purity, immune infiltration, or proliferation. For some phenotypes, it may not be clear whether the assumption holds. There may be spatial interactions and dependencies between molecular phenotypes that underlie a clinical phenotype. For example, neoadjuvant treatment drugs are designed to shrink regions of proliferative cells in breast cancer. However, it has also been observed that higher levels of tumor-infiltrating lympocytes are strongly associated with a higher rate of pathological complete response (pCR) [34]. SpacePhenotyper allows us to uncover such interactions between phenotypes. It is also possible that a phenotype depends on neighboring spatial regions. For example, tumor to stroma percentage is a marker of a poor prognosis in colorectal cancer [33]. This explains why SpacePhenotyper predicted high hazard not only on stroma/tumor boundary but also in a mostly stroma region (Figure 4). This underscores the importance of considering multiple molecular phenotypes for a proper interpretation of the relationship between local gene expression and clinical phenotypes.

In summary, we presented and validated a method for transferring phenotypic knowledge from bulk to the spatial domain. SpacePhenotyper outperformed other approaches on simulated data and produced biological results consistent with previously published findings. SpacePhenotyper represents an important novel computational tool for studying spatial transcriptomic data. Its framework is general, and we expect it to aid in the transfer of knowledge between other experimental techniques.

## Acknowledgment

B.A, E.H., Z.N.K, M.G.H., J.D.B., J.H., and T.M.P are supported by the NLM Intramural Research Program. D.L. and C.-P.D. is supported by the NCI Intramural Research Program. C.-P.D. is supported in part by FLEX Synergy Award from the NCI Center for Cancer Research. This research was supported in part by the Division of Intramural Research of the National Library of Medicine (NLM), National Institutes of Health (NIH) and by the National Cancer Institute (NCI), National Institutes of Health (NIH). The contributions of the NIH authors are considered Works of the United States Government. The findings and conclusions presented in this paper are those of the authors and do not necessarily reflect the views of the NIH or the U.S. Department of Health and Human Services. This work utilized the computational resources of the NIH HPC Biowulf cluster. (http://hpc.nih.gov) The results shown here are in part based upon data generated by the TCGA Research Network: https://www.cancer.gov/tcga.

## Supplementary Materials

## 1 Supplementary Methods

### 1.1 Data

#### TCGA gene expression data

The RSEM-estimated [35], FPKM-normalized [36], and batch-effect-corrected [37] TCGA gene expression data were downloaded from GitHub repository (https://github.com/mskcc/RNAseqDB) on May 20th, 2026. Requiring normal, cancer samples, their clinical information and availability of spatial transcriptomics data sets, Breast invasive carcinoma (BRCA) and Colon Adenocarcinoma (COAD) were obtained for the analysis and only patients whose clinical information is provided were kept for each cancer type. We excluded genes which are not significantly expressed in tumor, and the expression level of a gene is assumed as significant if 80% or more of its expression values over samples are greater than a threshold (see Fig. S2). The threshold is described by the minimum of absolute skew of normal distribution. We then selected the genes that are differentially expressed between normal and tumor samples (Benjamini Hochberg q-value ≤0.01), and that are also highly variable in the tumor samples. The variability cutoff of genes is decided based on minimum skew of variation coefficients (see Fig. S2, for more details about the processing of bulk gene expression data). The summary of the bulk gene expression cohorts is provided in Table S1.

#### Gene expression and Residual Cancer Burden (RCB)

Transcriptomic profiles of the pretreatment breast tumor biopsies from 168 patients representing all major breast cancer subtypes. FPKM normalization was performed to adjusts raw read counts to account for major technical biases that can distort gene expression measurements.

For the quantity measure of patients’ response to treatment, neoadjuvant chemotherapy was administered for a median of 18 weeks in 145 cases, primarily using sequential taxane and anthracycline regimens. Of the remaining patients, 22 received taxane-based treatment, while one received an anthracycline combined with cyclophosphamide. Two patients received only one cycle of chemotherapy because of drug-related toxicities. Patients with HER2-positive tumors (n = 65) received a median of three cycles of anti-HER2 therapy in combination with a taxane. Treatment response was evaluated after treatment using the residual cancer burden (RCB) scores.

#### Spatial transcriptomics data

The spatial transcriptomics data were downloaded from the 10X Genomics data set in which the samples were acquired from the following hyperlinks: Breast 1.1.0 (https://www.10xgenomics.com/datasets/human-breast-cancer-block-a-section-1-1-standard-1-1-0), Breast 1.3.0 (https://www.10xgenomics.com/datasets/human-breast-cancer-visium-fresh-frozen-whole-transcriptome-1-standard), Breast 2.0.0 (https://www.10xgenomics.com/products/xenium-in-situ/preview-dataset-human-breast), Colorectal 3.0.0 (https://cf.10xgenomics.com/samples/spatial-exp/3.0.0/Visium_HD_Human_Colon_Cancer/Visium_HD_Human_Colon_Cancer_web_summary.html)

### 1.2 Data simulation

#### Starting data

Human Lung Cell Atlas [38] single cell data was downloaded from synapse.org (accession ID: syn21043647, https://doi.org/10.7303/syn21041850). Read counts were obtained through SmartSeq-2 for 58683 transcripts in 9409 barcoded cells that are each annotated by one of 44 distinct cell types.

#### Preprocessing

From the single cell read count matrix, we remove all cells that do not belong to the following 3 cell types, leaving 551 epithelial Ciliated cells, 129 immune B cells, and 346 stromal Airway Smooth Muscle cells. Additionally, we restrict ourselves to transcripts representing genes from the TCGA-LUAD bulk expression data (downloaded from the Genomic Data Commons Data Portal (https://portal.gdc.cancer.gov/) on June 5th, 2020), shrinking the single cell read matrix down to 18408 genes and 1026 cells. We remove all genes whose median count is not > 0 in at least one cell type, resulting in 3273 genes remaining. Mitochondrial genes are also removed, resulting in 3257 remaining genes.

#### Synthetic SRT data

A 2D slice of 50 × 50 spatial locations was simulated. For each spot, a total number of all cells was selected uniformly at random to be between 4 and 10. The spots were organized into 4 regions displayed on the left side of Fig. S1, according to the following rules:

1. For the spots in the red region, 80% of their cells are Ciliated, and 20% of their cells are stromal.
2. For the spots in the blue region, 80% of their cells are B cells, and 20% of their cells are stromal.
3. For the spots in the purple region, 40% of their cells are Ciliated, another 40% of their cells are B cells, and the remaining 20% cells are stromal.
4. For the spots in the gray region, 100% of their cells are stromal.

Having chosen the number of cells of each type in each spot, we draw that many cells of corresponding types from the single cell data uniformly at random, and we add up gene read counts across all the selected cells for each gene as that spot’s final gene expression. The right side of Fig. S1 shows the resulting phenotype quantity heatmap for all the spots.

**Figure S1:**
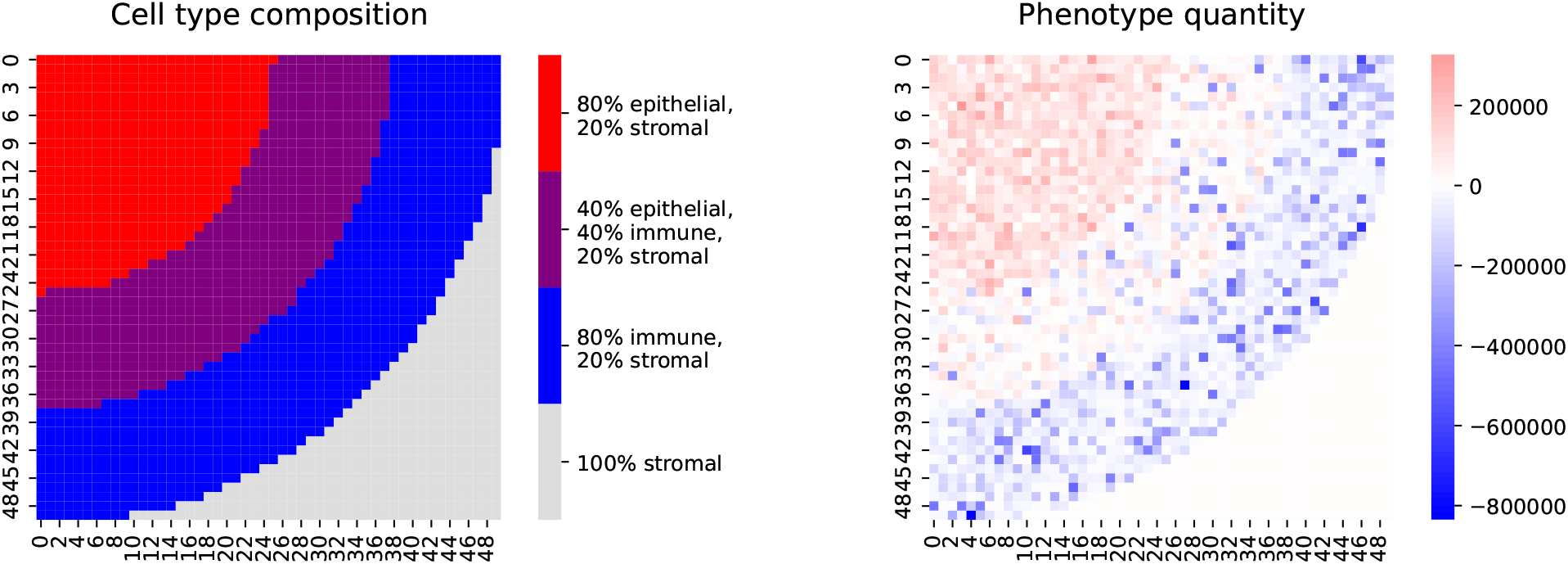
Synthetic SRT regions. The left side plot shows the cell type composition of the 4 distinct regions. The right side plot shows the phenotype quantities of all spots.

#### Synthetic Bulk data

We simulate 200 bulk patients and draw the number of cells of all 3 types for each patient from an exponential distribution with 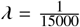, so that the average cell count for each cell type of a bulk patient is 15000 and the total number of cells in a bulk patient sample is on average 45000 (in line with the expected number of cells in a bulk-sequenced sample). Having chosen the number of cells of each type in each patient, we draw that many cells of corresponding types from the preprocessed single cell data uniformly at random, and we add up gene read counts across all the selected cells for each gene as that patient’s final bulk gene expression.

#### Gene-to-phenotype relation

From the 3257 genes obtained through preprocessing, we identify those that are expressed in all 3 cell types, and we sort them in descending order by their coefficient of variation 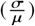. Starting from the top of this ordered list of genes, we select 10 genes for each cell type, such that the expression of those 10 genes is involved in determining the phenotype quantity in cells of the corresponding type. Thus, genes MALAT1, ACTG1, GPX3, TUBA1A, INMT, MT2A, KRT8, FOS, CSRP1, DEPP1 are selected to contribute positively to the simulated phenotype only in the Ciliated cells in which their collective expression is at least 1.5x the sum of their average expression over all 1026 cells (so that not all Ciliated cells contribute to the phenotype); let us label the sum of expression of these genes in all the cells that qualify as *sumA*. Genes TACSTD2, HSP90AA1, C9orf24, CALM1, DUSP1, CLU, TMSB4X, GSTP1, UCP2, GSN contribute negatively to the simulated phenotype only in B cells (no matter their expression level in a particular cell); let us label the sum of expression of these genes in all B cells as *sumB*. Stromal cells are left neutral with respect to the phenotype, and no genes expressed in them contribute to the bulk or spot phenotype quantity. For a particular bulk patient’s or SRT spot’s *sumA* and *sumB* values (calculated across all Ciliated and B cells in that patient or spot), the quantity of the simulated phenotype is calculated as *sumA* – *α* · *sumB*. The value of *α* was selected in order for the split of bulk patients with positive and negative phenotype quantities to be roughly half-half, resulting in *α* = 8.0.

## 2 Supplementary Tables

**Table S1:** Summary of the bulk gene expression cohorts, including the numbers of significantly expressed genes (Figure S2 A), differentially expressed genes (*q* < 0.01), highly variable genes (Figure S2 B), and patients.

| Cancer | Significantly<br>expressed genes | Differential<br>expressed genes | Highly<br>variable genes | Patients |
|---|---|---|---|---|
| TCGA BRCA | 13703 | 9445 | 3943 | 966 |
| TCGA COAD | 13675 | 8398 | 4532 | 279 |
| TransNEO BRCA | 4315 |  | 1677 | 162 |

## 3 Supplementary Figures

**Figure S2:**
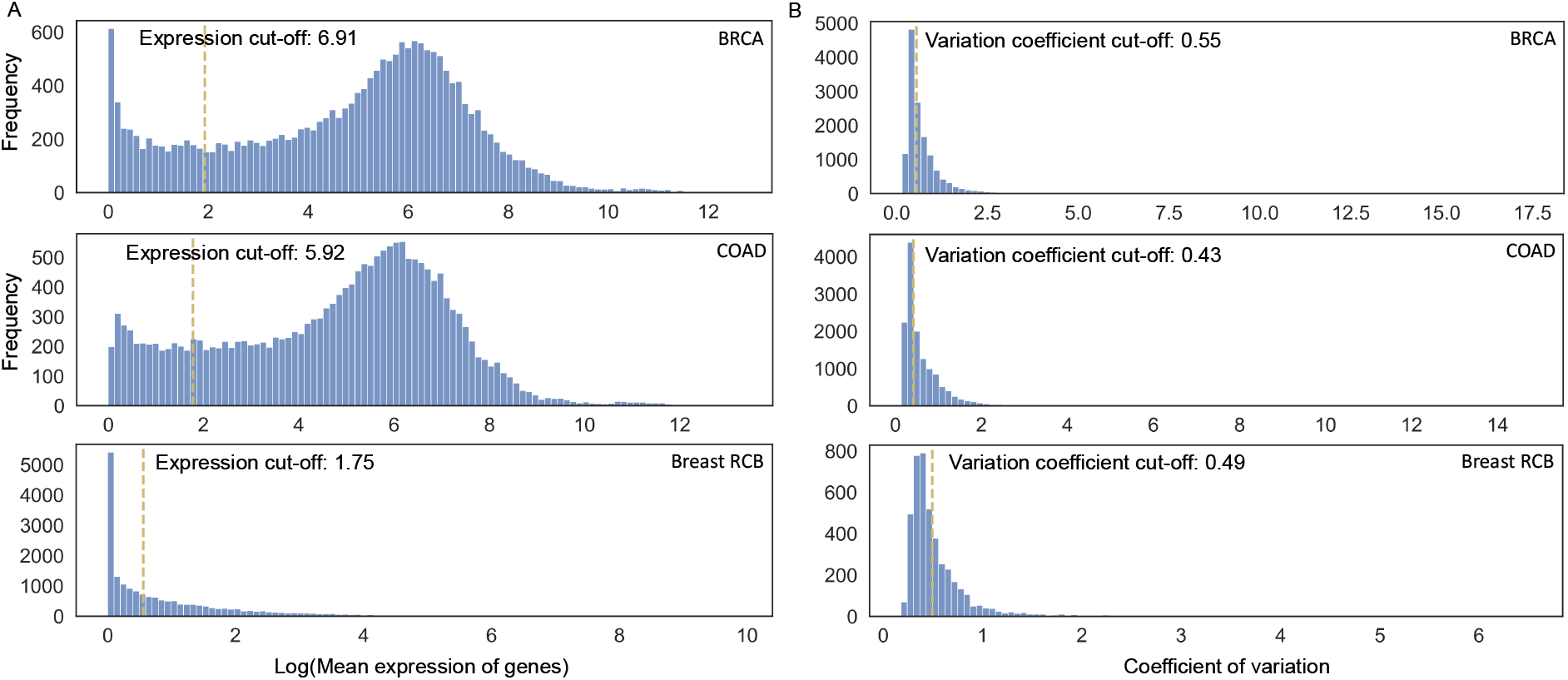
(A) Gene expression values exhibit a negative skew, with a substantial proportion of genes showing very low expression levels, potentially reflecting transcriptional inactivity. Assuming that the expression levels of actively expressed genes follow a normal distribution, we excluded lowly expressed genes using a cutoff determined by minimizing the absolute skewness of the mean expression distribution. (B) The distribution of gene coefficients of variation (CV) is positively skewed, indicating that a subset of genes exhibits high variability. Assuming that non-variable genes follow a normal distribution, we determined the cutoff for identifying highly variable genes by minimizing the absolute skewness of the CV distribution.

**Figure S3:**
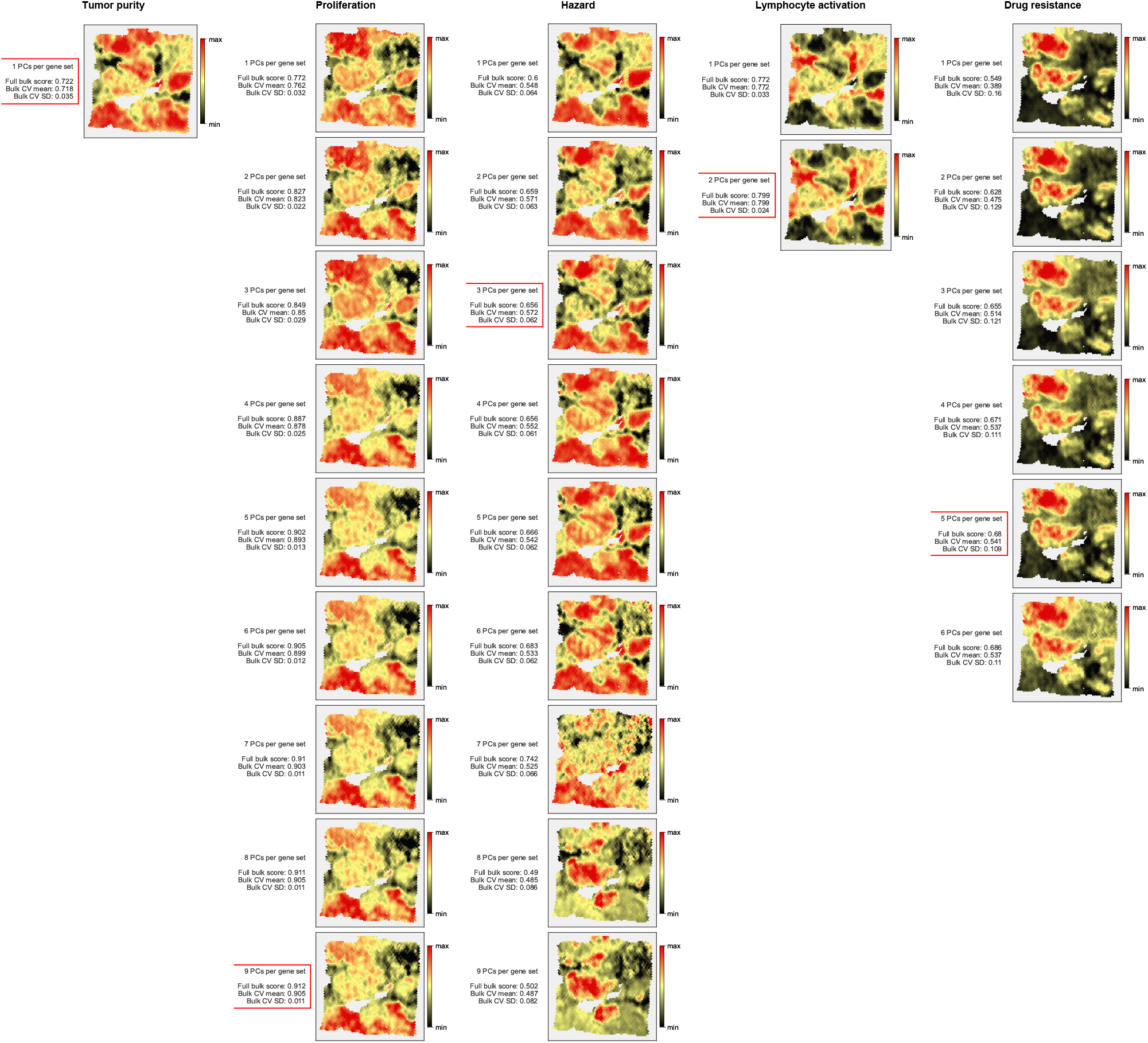
Predictions on Breast-1.1.0 with increasing numbers of selected principal directions. 10% genes with the highest absolute association with the phenotype were used. The red outline indicates the predictor with the best mean cross-validation score.

**Figure S4:**
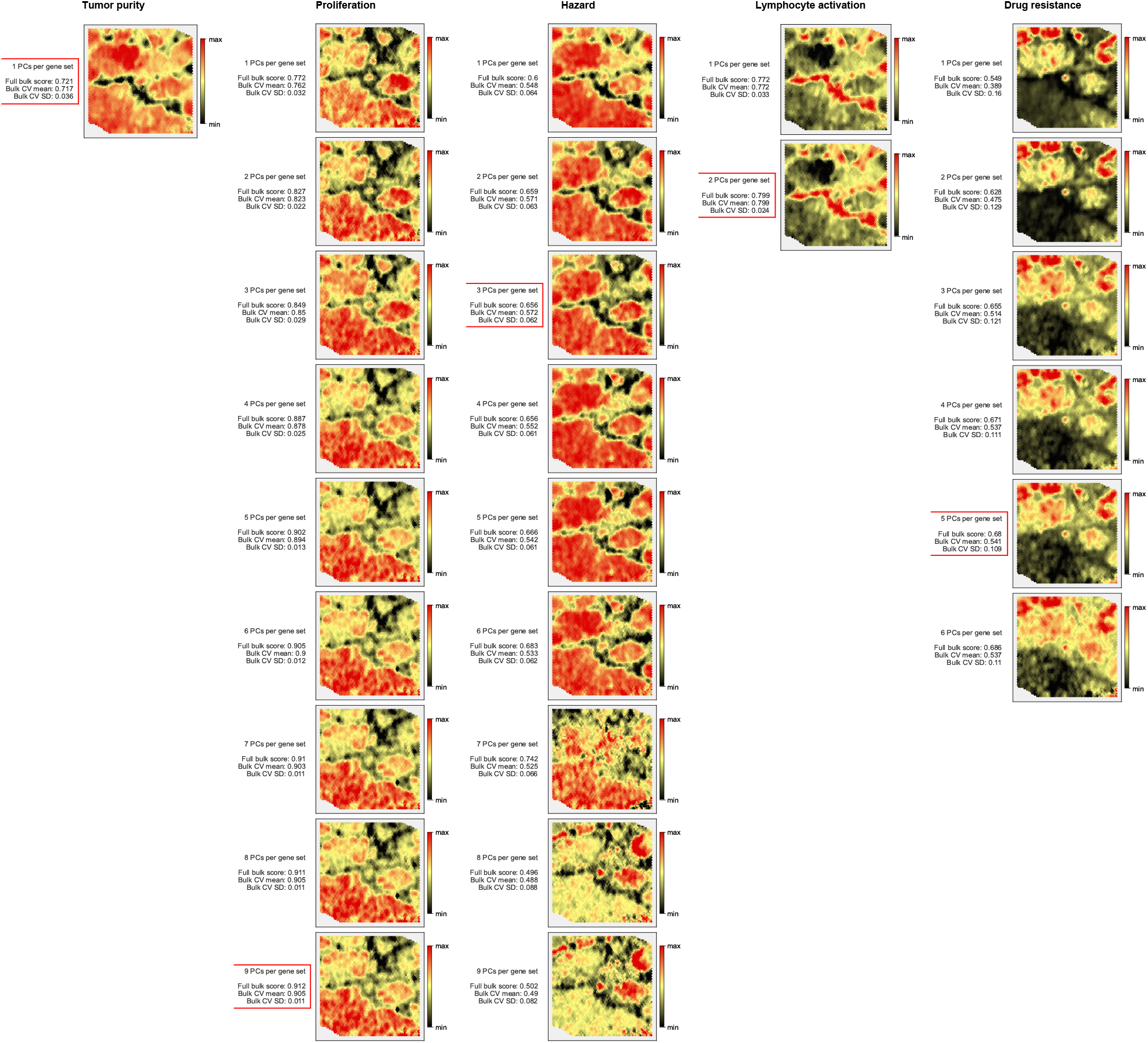
Predictions on Breast-1.3.0 with increasing numbers of selected principal directions. 10% genes with the highest absolute association with the phenotype were used. The red outline indicates the predictor with the best mean cross-validation score.

**Figure S5:**
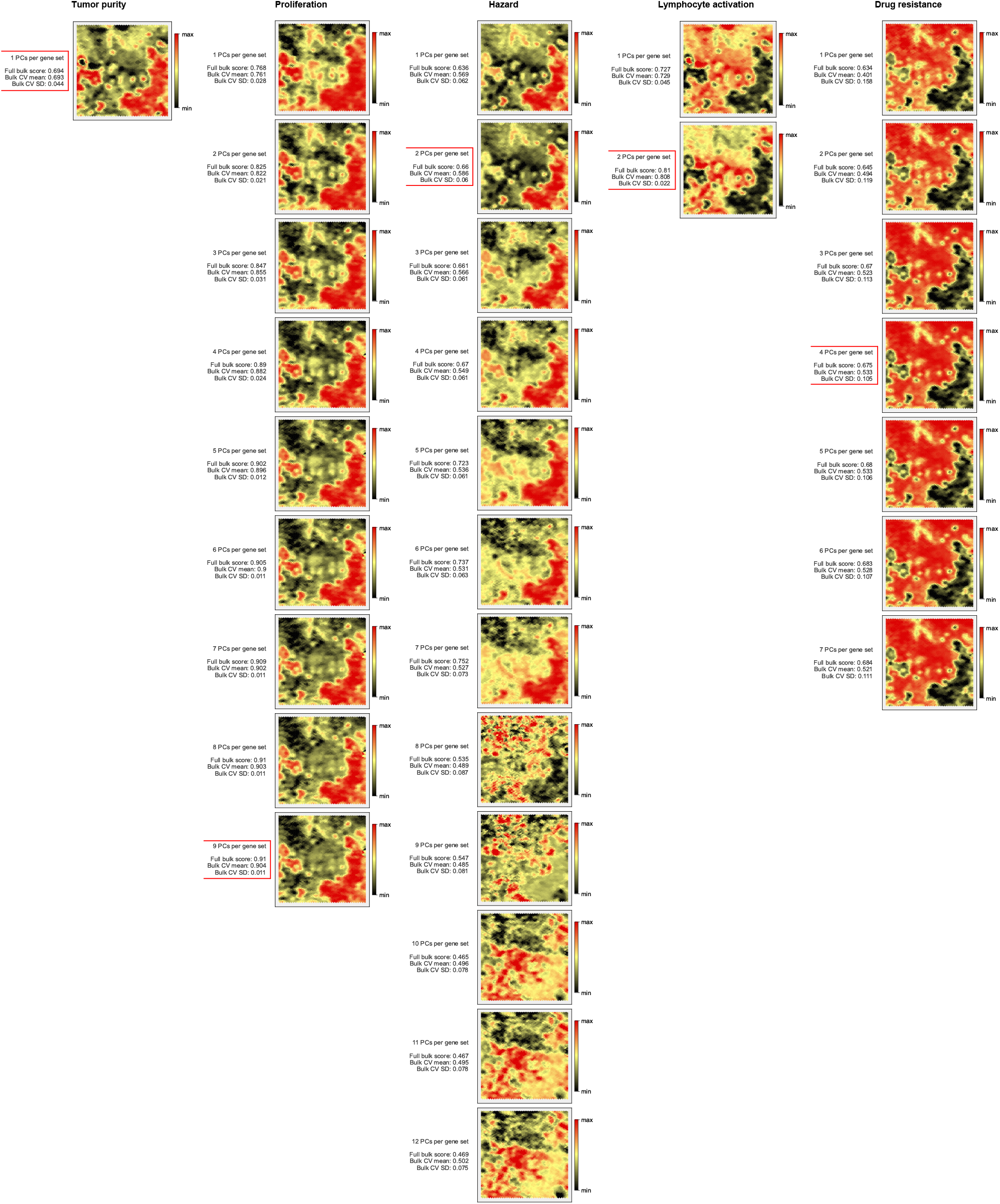
Predictions on Breast-2.0.0 with increasing numbers of selected principal directions. 10% genes with the highest absolute association with the phenotype were used. The red outline indicates the predictor with the best mean cross-validation score.

**Figure S6:**
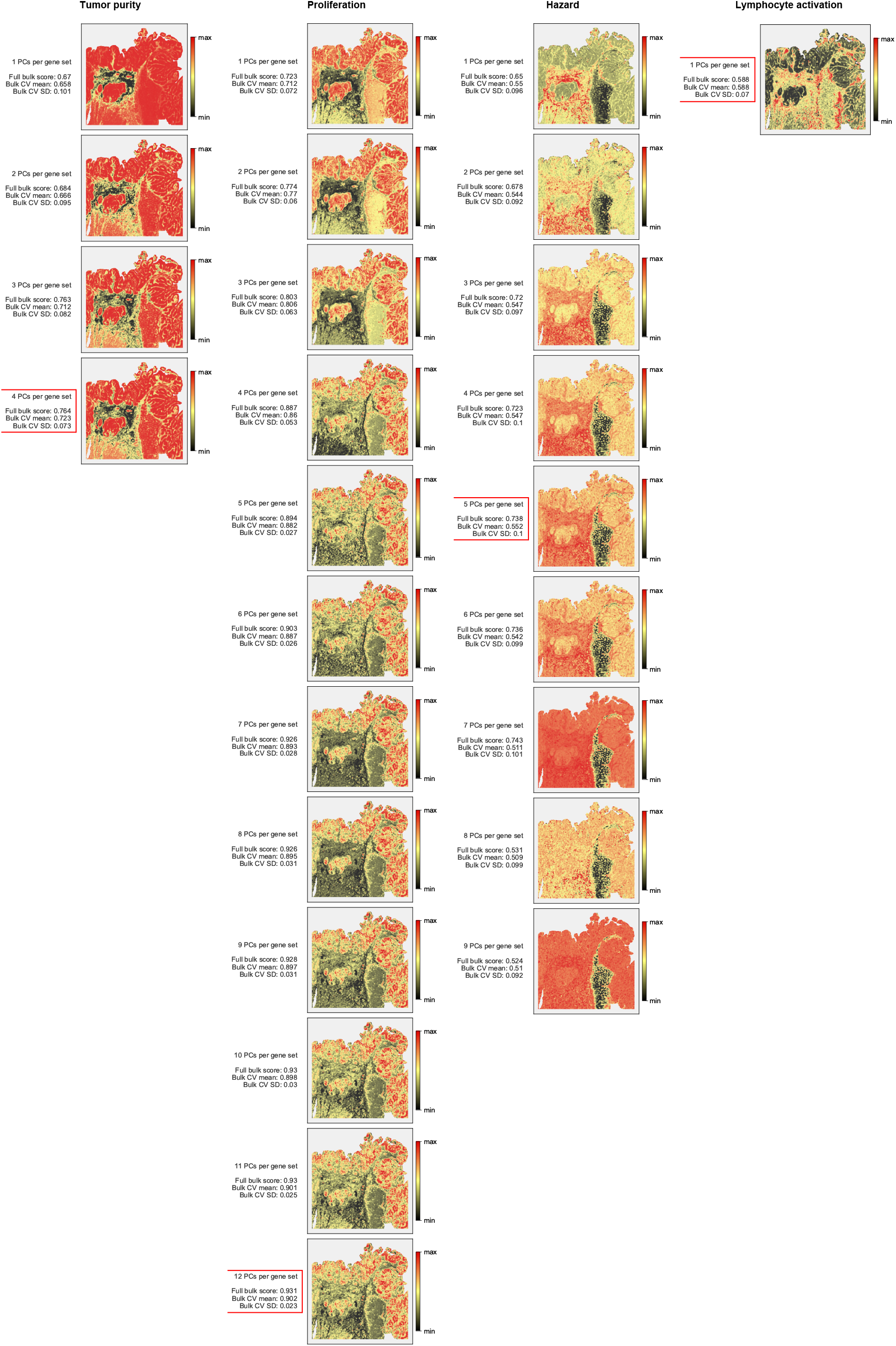
Predictions on Colorectal-3.0.0 with increasing numbers of selected principal directions. 10% genes with the highest absolute association with the phenotype were used. The red outline indicates the predictor with the best mean cross-validation score.

**Figure S7:**
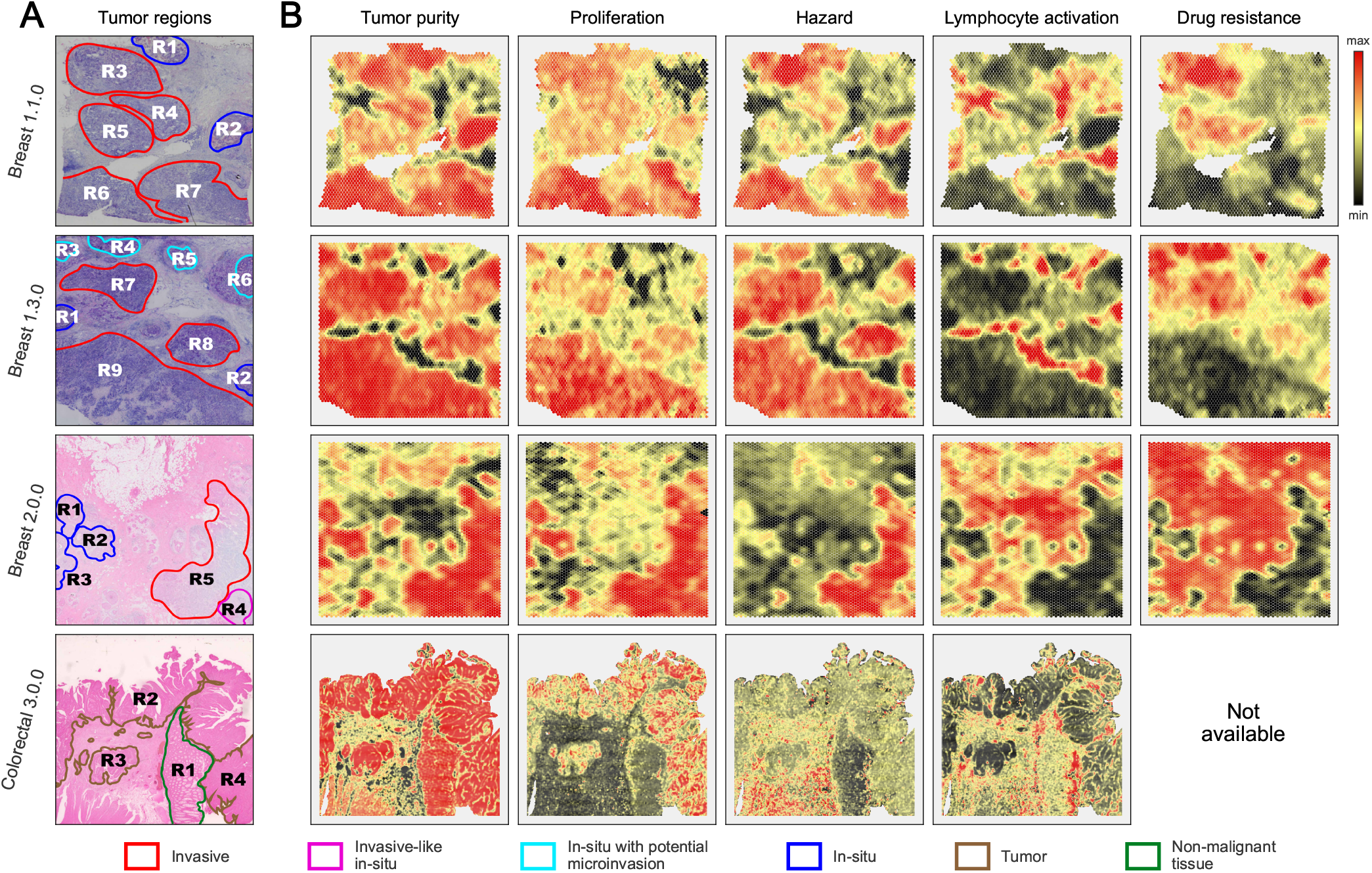
Phenotype predictions with 5% genes with the highest absolute association with the phenotype

**Figure S8:**
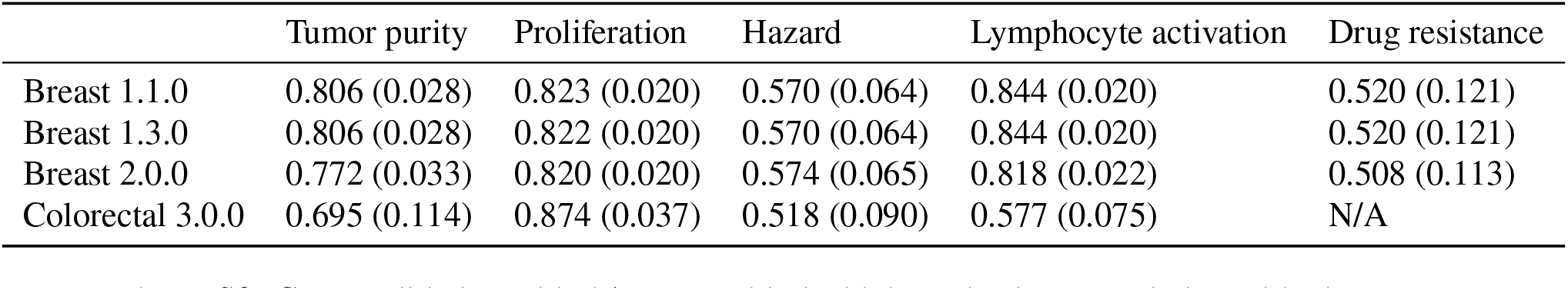
Cross-validation with 5% genes with the highest absolute association with phenotype

**Figure S9:**
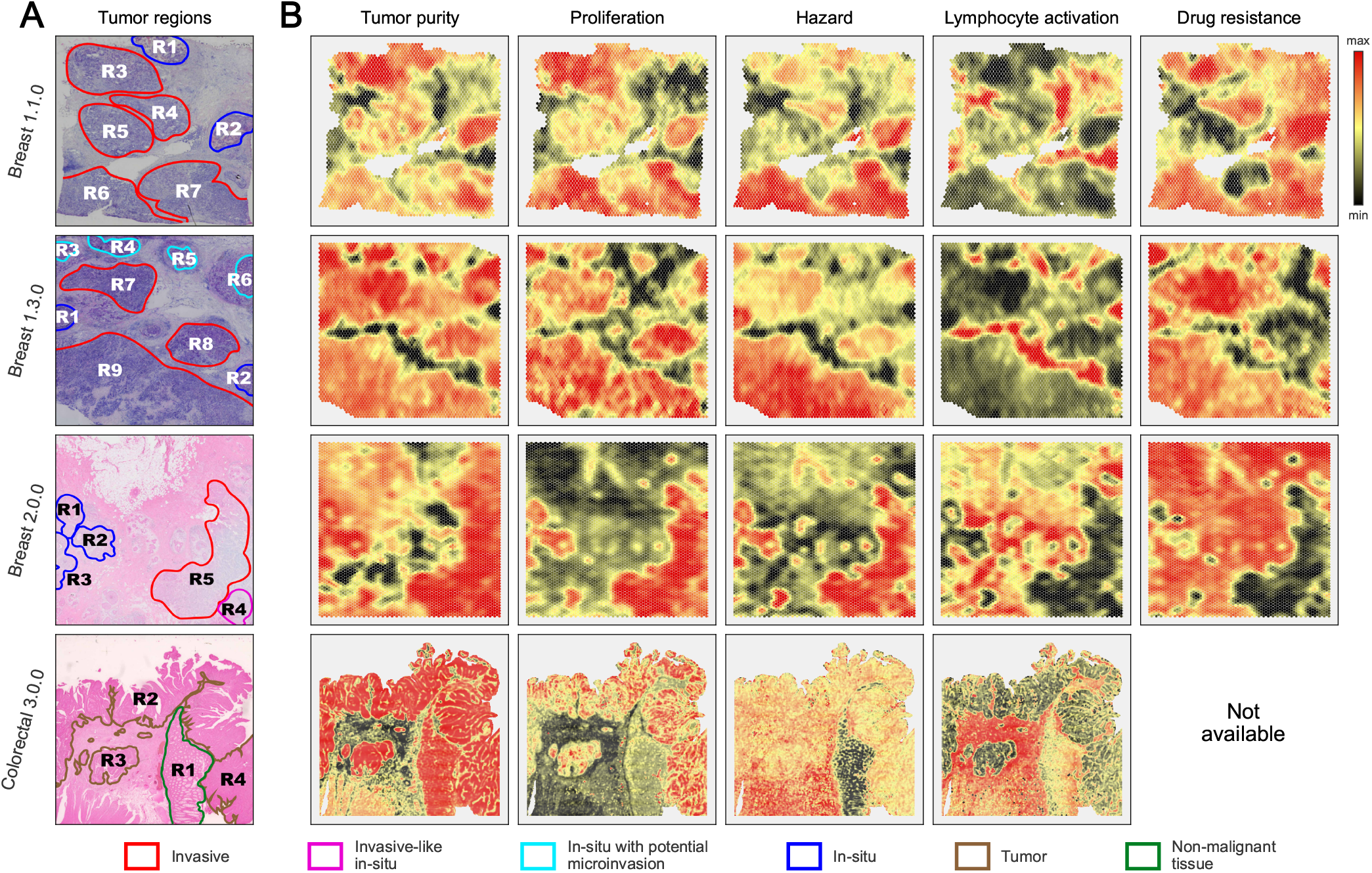
Phenotype predictions with 20% genes with the highest absolute association with the phenotype

**Figure S10:**
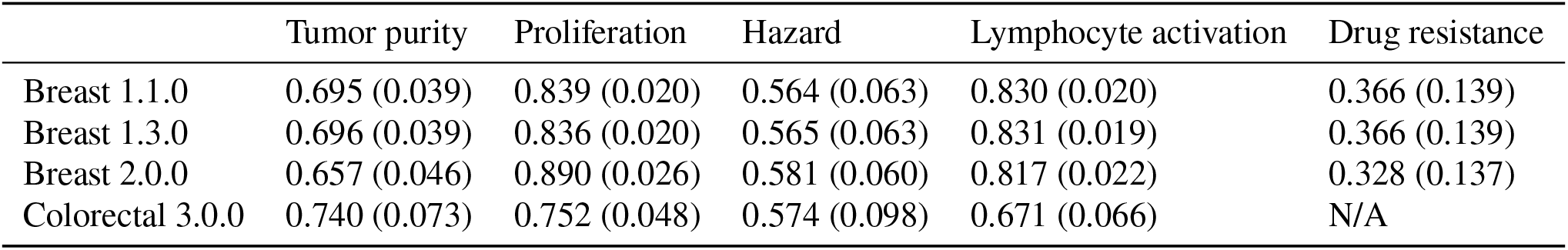
Cross-validation with 20% genes with the highest absolute association with phenotype

## Footnotes

1 See Supplementary Figures S7 to S10 for results with alternative percentage choices.

## References

[1] K. E. de Visser and J. A. Joyce. The evolving tumor microenvironment: From cancer initiation to metastatic outgrowth. Cancer Cell, 41(3):374–403, Mar 2023.

[2] B. Arneth. Tumor Microenvironment. Medicina (Kaunas), 56(1), Dec 2019.

[3] P. L. hl, F. n, S. Vickovic, A. Lundmark, J. F. Navarro, J. Magnusson, S. Giacomello, M. Asp, J. O. Westholm, M. Huss, A. Mollbrink, S. Linnarsson, S. Codeluppi, Å. Borg, F. n, P. I. Costea, P. n, J. Mulder, O. Bergmann, J. Lundeberg, and J. n. Visualization and analysis of gene expression in tissue sections by spatial transcriptomics. Science, 353(6294):78–82, Jul 2016.

[4] M. Cheng, Y. Jiang, J. Xu, A. A. Mentis, S. Wang, H. Zheng, S. K. Sahu, L. Liu, and X. Xu. Spatially resolved transcriptomics: a comprehensive review of their technological advances, applications, and challenges. J Genet Genomics, 50(9):625–640, Sep 2023.

[5] V. Marx. Method of the Year: spatially resolved transcriptomics. Nat Methods, 18(1):9–14, Jan 2021.

[6] M. Asp, J. hle, and J. Lundeberg. Spatially Resolved Transcriptomes-Next Generation Tools for Tissue Exploration. Bioessays, 42(10):e1900221, Oct 2020.

[7] B. L. Walker, Z. Cang, H. Ren, E. Bourgain-Chang, and Q. Nie. Deciphering tissue structure and function using spatial transcriptomics. Commun Biol, 5(1):220, Mar 2022.

[8] Y. Ma and X. Zhou. Spatially informed cell-type deconvolution for spatial transcriptomics. Nat Biotechnol, 40(9):1349–1359, Sep 2022.

[9] Y. Li and Y. Luo. STdGCN: spatial transcriptomic cell-type deconvolution using graph convolutional networks. Genome Biol, 25(1):206, Aug 2024.

[10] Z. Liu, D. Wu, W. Zhai, and L. Ma. SONAR enables cell type deconvolution with spatially weighted Poisson-Gamma model for spatial transcriptomics. Nat Commun, 14(1):4727, Aug 2023.

[11] K. Coleman, J. Hu, A. Schroeder, E. B. Lee, and M. Li. SpaDecon: cell-type deconvolution in spatial transcriptomics with semi-supervised learning. Commun Biol, 6(1):378, Apr 2023.

[12] Z. Zhou, Y. Zhong, Z. Zhang, and X. Ren. Spatial transcriptomics deconvolution at single-cell resolution using Redeconve. Nat Commun, 14(1):7930, Dec 2023.

[13] B. Ru, J. Huang, Y. Zhang, K. Aldape, and P. Jiang. Estimation of cell lineages in tumors from spatial transcriptomics data. Nat Commun, 14(1):568, Feb 2023.

[14] H. Sarkar, U. Chitra, J. Gold, and B. J. Raphael. A count-based model for delineating cell-cell interactions in spatial transcriptomics data. Bioinformatics, 40(Suppl 1):i481–i489, Jun 2024.

[15] Z. Cang, Y. Zhao, A. A. Almet, A. Stabell, R. Ramos, M. V. Plikus, S. X. Atwood, and Q. Nie. Screening cell-cell communication in spatial transcriptomics via collective optimal transport. Nat Methods, 20(2):218–228, Feb 2023.

[16] J. Zhu, Y. Wang, W. Y. Chang, A. Malewska, F. Napolitano, J. C. Gahan, N. Unni, M. Zhao, R. Yuan, F. Wu, L. Yue, L. Guo, Z. Zhao, D. Z. Chen, R. Hannan, S. Zhang, G. Xiao, P. Mu, A. B. Hanker, D. Strand, C. L. Arteaga, N. Desai, X. Wang, Y. Xie, and T. Wang. Mapping cellular interactions from spatially resolved transcriptomics data. Nat Methods, 21(10):1830–1842, Oct 2024.

[17] Zhaoyang Liu, Dongqing Sun, and Chenfei Wang. Evaluation of cell-cell interaction methods by integrating single-cell rna sequencing data with spatial information. Genome Biology, 23(1):218, 2022.

[18] K. Yoshihara, M. Shahmoradgoli, E. Martínez, R. Vegesna, H. Kim, W. Torres-Garcia, V. Treviño, H. Shen, P. W. Laird, D. A. Levine, et al. Inferring tumour purity and stromal and immune cell admixture from expression data. Nature communications, 4(1):2612, Oct 2013.

[19] M. Sibai, S. Cervilla, D. Grases, E. Musulen, R. Lazcano, CK. Mo, V. Davalos, A. Fortian, A. Bernat, M. Romeo, et al. The spatial landscape of cancer hallmarks reveals patterns of tumor ecological dynamics and drug sensitivity. Cell Reports, 44(2), Feb 2025.

[20] Bryan He, Matthew Thomson, Meena Subramaniam, Richard Perez, Chun Jimmie Ye, and James Zou. Cloudpred: Predicting patient phenotypes from single-cell rna-seq. In PACIFIC SYMPOSIUM ON BIOCOMPUTING 2022, pages 337–348. World Scientific, 2021.

[21] Eldad D. Shulman, Emma M. Campagnolo, Roshan Lodha, Youngmin Chung, Amos Stemmer, Thomas Cantore, Beibei Ru, Tian-Gen Chang, Sumona Biswas, Saugato Rahman Dhruba, Sumeet Patiyal, Sushant Patkar, Andrew Wang, Ranjan K. Barman, Chuhan Wang, Rohit Paul, Sarath Chandra Kalisetty, Tom Hu, MacLean P. Nasrallah, Ellis Patrick, Jean Yang, Yuan Yuan, Karine Sargsyan, Amy Plotkin, Padma Sheila Rajagopal, Stephen-John Sammut, Stanley Lipkowitz, Peng Jiang, Carlos Caldas, Simon R.V. Knott, Kenneth Aldape, Joo Sang Lee, Danh-Tai Hoang, and Eytan Ruppin. Ai-predicted spatial transcriptomics unlocks breast cancer biomarkers from pathology. Cell, 189(14):4225–4240.e25, July 2026.

[22] Xiaojie Cheng, Chen Tang, Kejing Dong, Yuzhou You, Xueying Zhao, Bin Duan, Shaoqi Chen, Guohui Chuai, Zhenbo Zhang, and Qi Liu. Spalinker identifies phenotype-associated spatial tumor microenvironment features by integrating bulk and spatial sequencing data. Cell Genomics, 5(7):100893, July 2025.

[23] Bin Duan, Xiaojie Cheng, and Hua Zou. Spapheno: linking spatial transcriptomics to clinical phenotypes with interpretable machine learning. Genome Medicine, 18(1), April 2026.

[24] DR. Cox. Regression models and life-tables. Journal of the Royal Statistical Society: Series B (Methodological), 34(2):187–202, 1972.

[25] W Fraser Symmans, Florentia Peintinger, Christos Hatzis, Radhika Rajan, Henry Kuerer, Vicente Valero, Lina Assad, Anna Poniecka, Bryan Hennessy, Marjorie Green, et al. Measurement of residual breast cancer burden to predict survival after neoadjuvant chemotherapy. Journal of Clinical Oncology, 25(28):4414–4422, 2007.

[26] D. Aran, M. Sirota, and A. J. Butte. Systematic pan-cancer analysis of tumour purity. Nat Commun, 6:8971, Dec 2015.

[27] A. Subramanian, P. Tamayo, V. K. Mootha, S. Mukherjee, B. L. Ebert, M. A. Gillette, A. Paulovich, S. L. Pomeroy, T. R. Golub, E. S. Lander, and J. P. Mesirov. Gene set enrichment analysis: a knowledge-based approach for interpreting genome-wide expression profiles. Proc Natl Acad Sci U S A, 102(43):15545–15550, Oct 2005.

[28] SJ. Sammut, M. Crispin-Ortuzar, SF. Chin, E. Provenzano, HA. Bardwell, W. Ma, W. Cope, A. Dariush, SJ. Dawson, JE. Abraham, et al. Multi-omic machine learning predictor of breast cancer therapy response. Nature, 601(7894):623–629, 2022.

[29] K. Dong and S. Zhang. Deciphering spatial domains from spatially resolved transcriptomics with an adaptive graph attention auto-encoder. Nat Commun, 13(1):1739, Apr 2022.

[30] S. Avesani, E. Viesi, L. ı, G. Motterle, V. Bonnici, M. Beccuti, R. Calogero, and R. Giugno. Stardust: improving spatial transcriptomics data analysis through space-aware modularity optimization-based clustering. Gigascience, 11, Aug 2022.

[31] A. Deshpande, M. Loth, D. N. Sidiropoulos, S. Zhang, L. Yuan, A. TF. Bell, Q. Zhu, WJ. Ho, C. Santa-Maria, D. M. Gilkes, et al. Uncovering the spatial landscape of molecular interactions within the tumor microenvironment through latent spaces. Cell systems, 14(4):285–301, Apr 2023.

[32] A. Janesick, R. Shelansky, AD. Gottscho, F. Wagner, SR. Williams, M. Rouault, G. Beliakoff, CA. Morrison, MF. Oliveira, JT Sicherman, et al. High resolution mapping of the tumor microenvironment using integrated single-cell, spatial and in situ analysis. Nature communications, 14(1):8353, 2023.

[33] J.H. Park, C.H. Richards, D.C. McMillan, P.G. Horgan, and C.S.D. Roxburgh. The relationship between tumour stroma percentage, the tumour microenvironment and survival in patients with primary operable colorectal cancer. Annals of Oncology, 25(3):644–651, March 2014.

[34] SJ Wood, Y Gao, JH Lee, J Chen, Q Wang, JL Meisel, and X Li. High tumor infiltrating lymphocytes are significantly associated with pathological complete response in triple negative breast cancer treated with neoadjuvant keynote-522 chemoimmunotherapy. Breast cancer research and treatment, 205(1):193–199, 2024.

[35] David J Bakewell and Ernst Wit. Weighted analysis of microarray gene expression using maximum-likelihood. Bioinformatics, 21(6):723–729, 2005.

[36] Cole Trapnell, Brian A Williams, Geo Pertea, Ali Mortazavi, Gordon Kwan, Marijke J Van Baren, Steven L Salzberg, Barbara J Wold, and Lior Pachter. Transcript assembly and quantification by RNA-Seq reveals unannotated transcripts and isoform switching during cell differentiation. Nature biotechnology, 28(5):511–515, 2010.

[37] Q. Wang, J. Armenia, C. Zhang, A. V. Penson, E. Reznik, L. Zhang, T. Minet, A. Ochoa, B. E. Gross, C. A. Iacobuzio-Donahue, D. Betel, B. S. Taylor, J. Gao, and N. Schultz. Unifying cancer and normal RNA sequencing data from different sources. Sci Data, 5:180061, Apr 2018.

[38] K. J. Travaglini, A. N. Nabhan, L. Penland, R. Sinha, A. Gillich, R. V. Sit, S. Chang, S. D. Conley, Y. Mori, J. Seita, G. J. Berry, J. B. Shrager, R. J. Metzger, C. S. Kuo, N. Neff, I. L. Weissman, S. R. Quake, and M. A. Krasnow. A molecular cell atlas of the human lung from single-cell RNA sequencing. Nature, 587(7835):619–625, 11 2020.

